# Cytoplasmic ribosomes on mitochondria alter the local membrane environment for protein import

**DOI:** 10.1101/2024.07.17.604013

**Authors:** Ya-Ting Chang, Benjamin A. Barad, Hamidreza Rahmani, Brian M. Zid, Danielle A. Grotjahn

## Abstract

Most of the mitochondria proteome is nuclear-encoded, synthesized by cytoplasmic ribosomes, and targeted to mitochondria post-translationally. However, a subset of mitochondrial-targeted proteins is imported co-translationally, although the molecular mechanisms governing this process remain unclear. We employ cellular cryo-electron tomography to visualize interactions between cytoplasmic ribosomes and mitochondria in *Saccharomyces cerevisiae*. We use surface morphometrics tools to identify a subset of ribosomes optimally oriented on mitochondrial membranes for protein import. This allows us to establish the first subtomogram average structure of a cytoplasmic ribosome on the surface of the mitochondria in the native cellular context, which showed three distinct connections with the outer mitochondrial membrane surrounding the peptide exit tunnel. Further, this analysis demonstrated that cytoplasmic ribosomes primed for mitochondrial protein import cluster on the outer mitochondrial membrane at sites of local constrictions of the outer and inner mitochondrial membrane. Overall, our study reveals the architecture and the spatial organization of cytoplasmic ribosomes at the mitochondrial surface, providing a native cellular context to define the mechanisms that mediate efficient mitochondrial co-translational protein import.

**SUMMARY:** Chang et al. present a membrane-guided approach for identifying a subset of cytoplasmic ribosomes oriented for protein import on the mitochondrial surface in *Saccharomyces cerevisiae* using cryo-electron tomography. They show that ribosomes cluster, make multiple contacts with, and induce local changes to the mitochondrial membrane ultrastructure at import sites.

## INTRODUCTION

Mitochondria are essential double-membrane organelles required for energy production, metabolism, and stress signaling in eukaryotic cells. Mitochondria contain their own genome and protein synthesis machinery (Pfanner, Warscheid et al. 2019). However, only a small percentage (<1%) of mitochondrial proteins are encoded by mitochondrial DNA (mtDNA), with the majority encoded by nuclear genes and synthesized by cytoplasmic ribosomes (Wiedemann and Pfanner 2017). These nuclear-encoded proteins are recognized by the receptors of mitochondrial import machinery through mitochondrial targeting sequences (MTS) localized at the N-terminus or internally within the polypeptide sequence. The MTS directs these proteins to the translocase of the outer membrane (TOM), which facilitates their translocation across the outer mitochondrial membrane (OMM). Proteins destined for the inner mitochondrial membrane (IMM) and matrix are transported through additional channels, including the translocase of the inner membrane 23 (TIM23) and translocase of the inner membrane 22 (TIM22) complexes (Chacinska, Koehler et al. 2009). The defective import of mitochondrial proteins through these pathways can lead to oxidative stress, neurodegenerative diseases, and metabolic diseases (MacKenzie and Payne 2007), underscoring the importance of mitochondrial protein import integrity for maintaining cellular health and viability.

The majority of nuclear-encoded mitochondrial proteins are synthesized by ribosomes in the cytosol and then post-translationally targeted to the mitochondria for protein import (Chacinska, Koehler et al. 2009, Avendaño-Monsalve, Ponce-Rojas et al. 2020). However, electron microscopy studies conducted nearly five decades ago detected cytoplasmic ribosomes in proximity to the outer mitochondrial membrane (Kellems, Allison et al. 1975), suggesting that some proteins may be co-translationally imported into this organelle. Consistent with this, later work showed that a population of mRNAs encoding mitochondrial proteins was enriched in the vicinity of mitochondria (Suissa and Schatz 1982, Marc, Margeot et al. 2002, Saint-Georges, Garcia et al. 2008, Tsuboi, Viana et al. 2020). Further, proximity ribosome profiling identified a subset of mitochondrial proteins, highly enriched for IMM proteins, that are subject to co-translational import (Williams, Jan et al. 2014).

While it is clear that a population of mitochondrial proteins are co-translationally imported, the structural and mechanistic basis of this process remains poorly understood. The nascent polypeptide-associated complex (NAC) has been shown to promote the interaction between ribosomes and mitochondria to stimulate mitochondrial protein import (George, Walsh et al. 2002, Lesnik, Cohen et al. 2014). OM14 on the OMM was further identified as a receptor for ribosome-NAC complex during co-translational import (Lesnik, Cohen et al. 2014). In addition to the interface between OM14 and NAC, the interaction between TOM complex and nascent peptide appears crucial for ribosome recruitment on OMM (Pfanner, Wiedemann et al. 2004, Schulz, Schendzielorz et al. 2015, Gold, Chroscicki et al. 2017, Wang, Chen et al. 2020). Although these studies illustrate the occurrence of co-translational import in mitochondria, the function, regulation, and molecular composition of co-translational import in native cells remains to be defined.

Cryo-electron tomography (cryo-ET) is an imaging technique that produces detailed three-dimensional (3D) reconstructions of organelles and macromolecules in their native cellular environment. When combined with subtomogram averaging, cryo-ET can visualize endogenous macromolecular protein complexes at subnanometer resolution (Young and Villa 2023). These approaches have been applied to reveal the recognition and import mechanisms of co-translating ribosomes docked on the endoplasmic reticulum (ER) membrane (Pfeffer, Brandt et al. 2012, Pfeffer, Burbaum et al. 2015, Gemmer, Chaillet et al. 2023). However, in contrast to the highly abundant co-translation events observed on the ER membrane, mitochondrial co-translation is comparatively rare and, therefore, much more difficult to structurally characterize using these methods owing to the low numbers of ribosomes localized to the OMM (Gold, Chroscicki et al. 2017). Therefore, the regulatory mechanisms that stabilize this association to promote efficient co-translational import have yet to be revealed.

Here, we used cryo-focused ion beam (cryo-FIB) milling and cryo-ET to capture endogenous associations between cytoplasmic ribosomes and mitochondrial membranes in *Saccharomyces cerevisiae* (*S. cerevisiae*). We leveraged our previously developed Surface Morphometrics pipeline (Barad, Medina et al. 2023) to identify a subset of cytoplasmic ribosomes with their peptide exit tunnel optimally oriented for co-translational import on the OMM. Subtomogram averaging analysis of these mitochondrially-associated ribosomes revealed the first structure of mitochondria-associated ribosomes oriented for protein import and identified multiple contacts between the ribosome and OMM formed around the peptide exit tunnel. We show that these ribosomes tend to cluster on the OMM in an arrangement suggestive of polysome formation. Surprisingly, we observe a decrease in the OMM-IMM distance locally at co-translational import sites, suggesting that these membrane regions may be optimally remodeled for efficient protein import. Our study provides insight into the previously uncharacterized structural interactions of cytoplasmic ribosomes at mitochondrial membranes that facilitate mitochondrial protein import in cells.

## RESULTS AND DISCUSSION

### Cellular cryo-electron tomography captures cytoplasmic ribosomes surrounding mitochondria in native cellular conditions

Given that mitochondrial protein co-translation events are comparatively rare relative to the highly abundant and prevalent ER co-translation events, we sought to establish the optimal conditions for enriching mitochondrially-associated ribosomes in *S. cerevisiae*. Previous work demonstrated that mitochondrial mRNA localization increases in yeast grown in respiratory conditions relative to fermentative conditions (Tsuboi, Viana et al. 2020). This suggested that mitochondrial protein co-translation may similarly increase under these conditions. Additionally, treatment with translation elongation inhibitors such as cycloheximide (CHX) arrests ribosomes on the mitochondrial membrane and increases the number of mitochondrial-associated ribosomes that co-purify with mitochondria from *S. cerevisiae* (Gold, Chroscicki et al. 2017). Therefore, we grew yeast in fermentative or respiratory conditions and in the presence or absence of CHX prior to cellular fixation using standard vitrification procedures (**Figure 1A**). We used cryo-fluorescence microscopy to image vitrified cells on the electron microscopy grids and select targets based on quality, such as cell density and ice thickness (**Figure 1B**). We then targeted cell clumps for cryo-FIB-milling to generate thin (∼150-200 nm) lamella (**Figure 1C**). Next, we collected tilt series datasets of the milled lamella at a lower magnification (pixel size = 2.638) to enable a larger field of view for the cellular context. In all conditions, we observed individual ribosomes at the surface of mitochondria in our reconstructed 3D tomograms (**Figure 1D, Supplementary Figure 1A**). As expected, CHX treatment increased these associations in both growth conditions (**Supplementary Figure 1B**). Overall, this demonstrates that, consistent with *in vitro* work (Gold, Chroscicki et al. 2017), CHX treatment enriches cytoplasmic ribosomes surrounding mitochondria within the cellular context.

**Figure 1.**
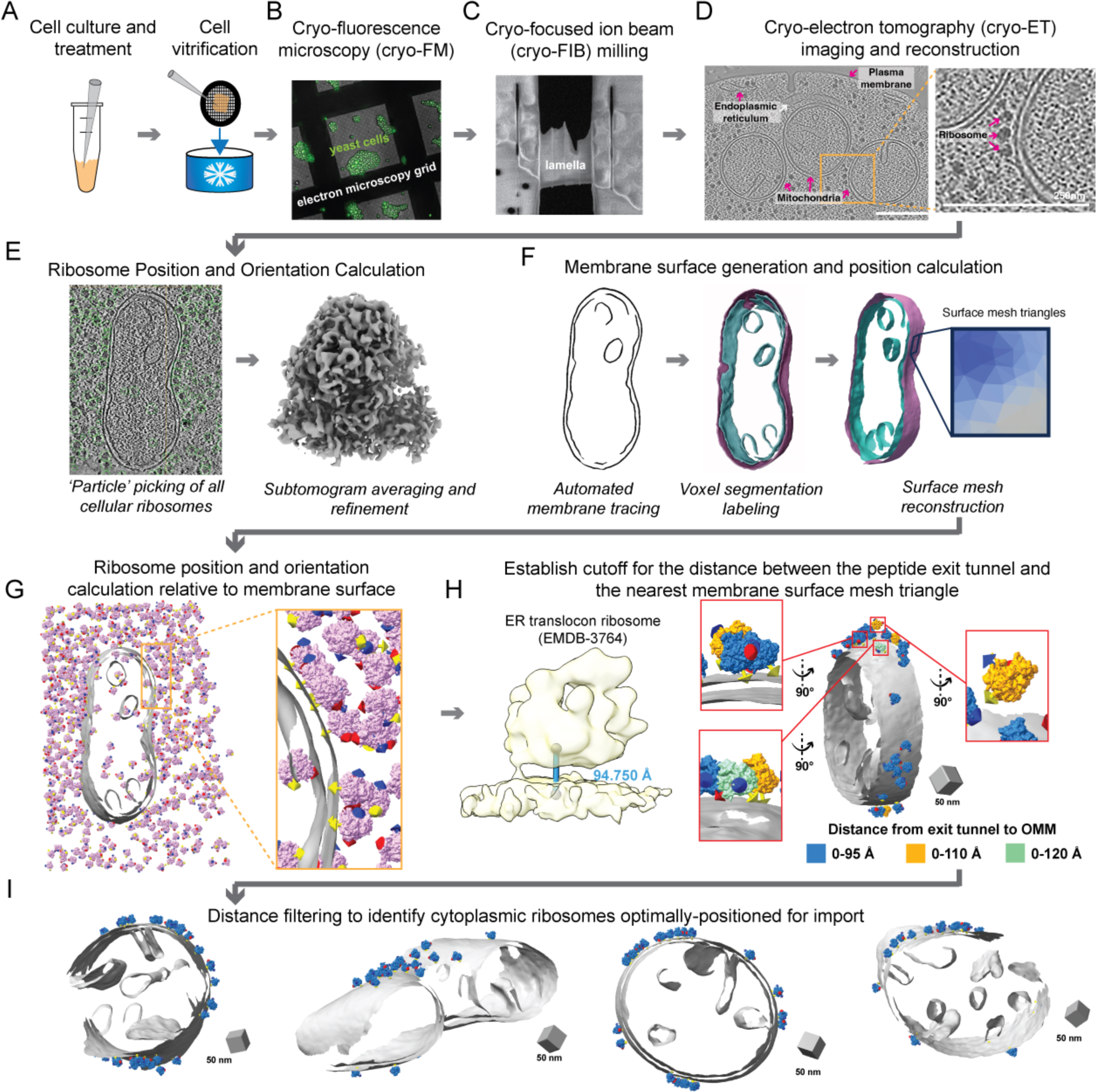
Cellular cryo-electron tomography imaging and processing workflow captures cytoplasmic ribosomes positioned for protein import on mitochondrial membranes. A. *Saccharomyces cerevisiae* (*S. cerevisiae*) yeast expressing TIM50-GFP are grown in respiratory or fermentative conditions and treated with vehicle or cycloheximide (CHX, 100 μg/mL) prior to deposition on electron microscopy grids (black mesh circle) and vitrification via plunge freezing. B. Vitrified yeast were imaged by cryo-fluorescence microscopy (cryo-FM) to assess sample quality, cell density, and ice thickness. C. Clumps of yeast were targeted for cryo-focused ion beam (cryo-FIB) milling to generate thin cellular sections (i.e., lamellae). D. Cellular lamella were imaged by standard cryo-electron tomography acquisition procedures to generate tilt series that were further processed to generate three-dimensional reconstructions (i.e., tomograms). Subcellular components such as mitochondria, the endoplasmic reticulum, the plasma membrane, and ribosomes are visible within the resulting tomograms. Scale bars = 250 nm E. Reconstructed tomograms were processed through ‘particle picking’ software, which identified the initial positions and orientations of all visible cellular ribosomes. The positions and orientations were refined using subtomogram averaging to produce a consensus 8 Å 80S ribosome structure. F. Mitochondrial membranes were traced, and separate three-dimensional voxel segmentations were generated for the outer and inner mitochondrial membranes (OMM and IMM, respectively). These voxel segmentations were converted to surface mesh reconstructions using the Surface Morphometrics (Barad, Medina et al. 2023) pipeline such that the location of the membrane is represented by the coordinate of each triangle within the mesh. G. The position and orientation of each ribosome relative to the OMM surface mesh reconstruction was calculated and rendered in the ArtiaX module of ChimeraX. The three-color arrows on ribosomes represent the Euler angle, with the yellow arrow representing the orientation of the ribosome peptide exit tunnel. H. The cutoff for identifying cytoplasmic ribosomes engaged in protein import on OMM was established by referring the distance between the peptide exit tunnel of ER-translocon ribosome and ER membrane. The optimal cutoff of the distance between exit tunnel and OMM was identified as 0-95 Å in ArtiaX as we started to observe the exit tunnel pointed away from OMM in the expanded cutoff, either 0-110 Å or 0-120 Å. I. Cytoplasmic ribosomes optimally positioned for protein import were identified as those with their exit tunnel closer than 95 Å from the OMM.

### Contextual surface morphometrics pipeline identifies cytoplasmic ribosomes optimally positioned for protein import on mitochondrial membranes

We set out to define the interactions between cytoplasmic ribosomes and the OMM likely responsible for mitochondrial protein co-translational import in the native cellular context using subtomogram averaging. The resolution attainable by these averaging approaches depends on several factors, including the magnification (i.e., pixel size) during acquisition, the signal-to-noise ratio, and the number of individual macromolecules (i.e., particles) present within the resulting tomographic data (Young and Villa 2023). With these parameters in mind, we collected an additional dataset of CHX-treated cells vitrified and milled into very thin (∼130 nm) lamella at higher magnification (pixel size = 1.6626). The resulting tomograms reveal the nanoscale architecture of cytoplasmic ribosomes in proximity to subcellular features such as mitochondria, the ER, and the plasma membrane (**Supplementary Figure 1C**). We developed a membrane-guided approach to identify the subset of ribosomes within the cellular milieu that is optimally positioned for mitochondrial protein import (**Figure 1E-H**). We computationally identified the position and initial orientations of all individual ribosomes within our dataset using automated 3D template matching software (Hrabe, Chen et al. 2012, Maurer, Siggel et al. 2024) (**Figure 1E)**. This resulted in the identification of 35,784 individual ribosomes from 91 tomograms. We further refined the initial positions and orientations using subtomogram averaging to produce an 8 Å map of the 80S ribosome (**Figure 1E, Supplementary Figure 1D**).

Next, we used our recently developed Surface Morphometrics pipeline (Barad, Medina et al. 2023) to generate surface mesh reconstructions from the voxel segmentations of mitochondrial membranes generated by automated tracing methods (Lamm, Zufferey et al. 2024) (**Figure 1F)**. Each membrane surface mesh consists of hundreds of thousands of individual triangles to represent the position and implicit geometry of mitochondrial membranes visible in the cellular tomograms. We then calculated the distance and orientation for each individual ribosome relative to the nearest surface mesh triangle on the OMM using Python-based scripts (**Figure 1G**). We reasoned that ribosomes engaged in mitochondrial protein import in cells would likely adopt an orientation similar to that of co-translating ribosomes on the ER (Pfeffer, Burbaum et al. 2015, Gold, Chroscicki et al. 2017) (EMD-3764) and on purified mitochondrial membranes (Gold, Chroscicki et al. 2017) (EMD-3762), with the peptide exit tunnel positioned within 95 Å from the membrane (**Figure 1H**). We, therefore, used this distance cut-off to identify the population of mitochondrially-associated cytoplasmic ribosomes likely engaged in co-translational protein import (n=1,076), which was confirmed through manual visual inspection (**Figure 1I, Supplementary Figure 2**). Overall, this demonstrates that only a subset of ribosomes is positioned optimally for import on the mitochondrial surface within the cellular context, consistent with previous work (Gold, Chroscicki et al. 2017).

### Multiple contacts form between the outer mitochondrial membrane and the subset of cytoplasmic ribosomes optimally positioned for protein import

Previous work using cryo-ET in combination with subtomogram averaging approaches has revealed that multiple contacts are formed between the cytoplasmic ribosome and the ER membrane to facilitate co-translational import into the ER lumen (Becker, Bhushan et al. 2009, Pfeffer, Brandt et al. 2012, Pfeffer, Burbaum et al. 2015, Jomaa, Gamerdinger et al. 2022, Gemmer, Chaillet et al. 2023, Jaskolowski, Jomaa et al. 2023). We wondered whether similar contact points are observed between cytoplasmic ribosomes and the OMM to facilitate protein import into mitochondria. We performed subtomogram averaging of the subset of ribosomes optimally positioned for mitochondrial co-translational import (**Figure 1I, Supplementary Figure 2**), which produced a final map resolved to 19 Å (**Figure 2, Supplementary Figure 3 & 4, Supplementary Movie 1**). In the resulting structure, we observe three contact points between the associated ribosome and OMM that surround the peptide exit tunnel (labeled 1, 2, 3 in **Figure 2, Supplementary Figure 3B**). To determine whether these contact points were specific to those optimally positioned for protein import, we averaged all ribosomes that were within 250 Å of the OMM but did not have their exit tunnel facing the OMM, which caused these three contact points to disappear (**Supplementary Fig. 3D**). Together, these data show multiple contacts are formed between the OMM and cytoplasmic ribosomes optimally positioned for protein import, suggesting that these connections likely stabilize co-translational protein import to the mitochondria.

**Figure 2.**
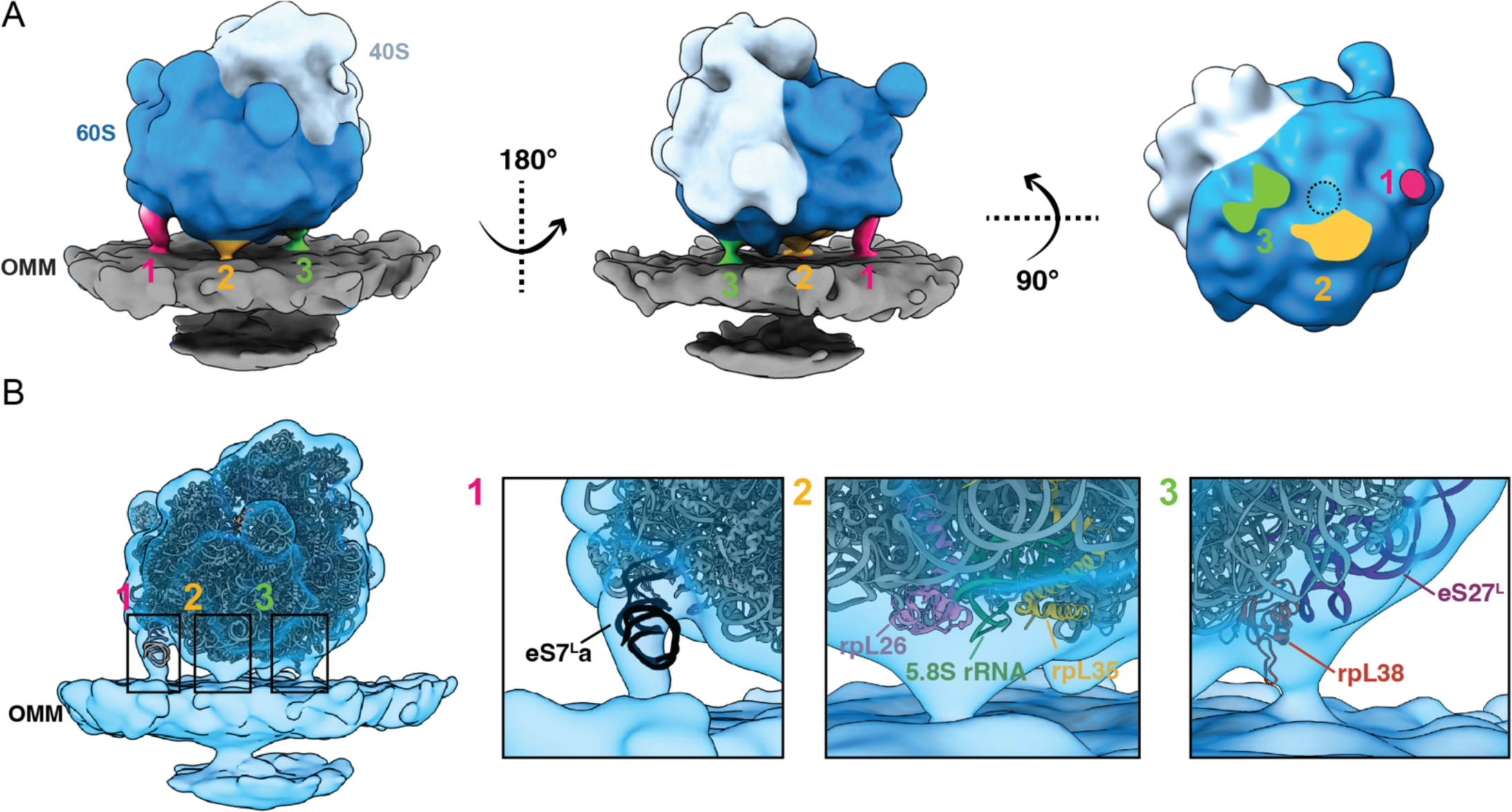
Three-dimensional subtomogram average of a cytoplasmic ribosome optimally positioned for protein import on the outer mitochondrial membrane (OMM). A. Three views of the subtomogram average of a cytoplasmic ribosome positioned with the exit tunnel on the 60S subunit (dark blue) facing the OMM (gray). Three connecting densities (labeled 1, 2, 3 in pink, orange, and green, respectively) are visible between the 60S and the OMM surrounding the peptide exit tunnel (dashed circle). B. The subtomogram average of the mitochondria-associated ribosome (blue transparent density) with a fitted atomic model of the *S. cerevisiae* 80S ribosome (PDB 4V6I). Boxed regions focus on the cryo-EM density of each of the three connections observed between the cytoplasmic ribosome and the OMM.

To further characterize these contacts, we rigid-body docked an atomic model of the 80S ribosome (PDB 4V6I) into the subtomogram average (**Figure 2B, Supplementary Figure 4**). The expansion segment of the eS7La of the 25S rRNA in the large subunit occupies the density corresponding to the connection labeled #1 in **Figure 2B**. This is consistent with previous subtomogram structures of ER-associated ribosomes from purified microsomes (EMDB-3764) (Gold, Chroscicki et al. 2017) (**Supplementary Figure 4A**), suggesting that a similar physical contact between this ribosomal RNA and the mitochondrial membrane may occur in protein import. Despite its prevalence in both organellar import systems across species, the functional role of this connection remains unknown. Future work is needed to determine how the rRNA attaches to membranes, either through direct lipid interaction or a protein receptor.

We observe a second connection (connection #2) between the cytoplasmic ribosome and the OMM formed immediately adjacent to the peptide exit tunnel (labeled 2 in **Figure 2**). At the current resolution of our subtomogram average, it is not feasible to unambiguously model polypeptide into this density to confirm its identity. However, several candidate molecules may be forming this connection. Given its proximity to the peptide exit tunnel, this connection may be formed by the nascent polypeptide chain itself. On the ribosome side, this connection is near ribosomal proteins rpL35, rpL26, and H5/6/7 from 5.8S rRNA (**Figure 2B)**. In previous structural work, these components were associated with the yeast Ssh1 complex (**Supplementary Figure 4B)** (EMDB-1667) (Becker, Bhushan et al. 2009), the homolog of the Sec61 import channel in the ER. The cytoplasmic loops of Ssh1 facilitate the docking and stabilization of cytoplasmic ribosomes on the ER membrane. These interactions promote the alignment of the peptide exit tunnel with the central pore of the channels for the import of the nascent chain across or through the membrane. In mitochondria, subunits of the TOM complex play an analogous role to the Ssh1/Sec61 complex in facilitating protein import (Aviram and Schuldiner 2017, Wiedemann and Pfanner 2017). Although the resolution of our subtomogram average is not sufficient to resolve any putative channel densities within the OMM, based on the positioning of connection #2 near the peptide exit tunnel, we propose that the TOM complex is likely situated directly underneath this connection on the OMM (**Supplementary Figure 4D**). The cytoplasmic portion of TOM20 subunit of the TOM complex has been shown to play a role in binding and stabilizing the incoming polypeptide during protein import (Ornelas, Bausewein et al. 2023). Interestingly, positioning an atomic model generated from a recently solved structure of the TOM complex by single-particle cryo-EM (PDB 8B4I) (Ornelas, Bausewein et al. 2023) in the OMM directly underneath the peptide exit tunnel shows that the globular, cytoplasmic-exposed portion of the TOM20 complex fits partially in our subtomogram average near the density corresponding to connection #2 (**Supplementary Figure 4D**), suggesting that it may be facilitating import at this region.

Several other protein biogenesis factors associate directly with the ribosome and engage with the nascent chain as it emerges from the peptide exit tunnel. This includes the NAC, which binds to nascent polypeptides as they emerge from the ribosomal peptide exit tunnel through contacts on the ribosome via rpL25/35 (Nyathi and Pool 2015, Deuerling, Gamerdinger et al. 2019). Previous work demonstrated that the NAC promotes the interaction between cytoplasmic ribosomes and the OMM through the outer membrane protein OM14 (Lesnik, Cohen et al. 2014); however, the exact structural interactions between the NAC and OM14 are unknown. In our subtomogram average structure, rpL35 is positioned near connection #2, suggesting that this density may correspond to the tethering between NAC and OM14 to promote mitochondrial co-translation. In support of this, connection #2 in our subtomogram average overlaps with the localization of the NAC when it has been displaced by the signal recognition particle (SRP) for translocation to the ER (Jomaa, Gamerdinger et al. 2022). This suggests that NAC may adopt a similar localization to mediate protein translocation in the OMM (**Supplementary Figure 4D**); however, future genetic, biochemical, and structural work is needed to determine the identity of this connection.

We observe a third connection between ribosome and OMM near the expansion segment eS27L of 25S rRNA and the ribosomal protein rpL38 (labeled 3 in **Figure 2**). A similar contact between the rRNA expansion segment, rpL38, and TRAP complex has been reported in the ER-associated ribosome (EMDB-3068, EMDB-16232, EMDB-15884) (Pfeffer, Burbaum et al. 2015, Gemmer, Chaillet et al. 2023, Jaskolowski, Jomaa et al. 2023) (**Supplementary Figure 4C**). The TRAP complex interacts with cytoplasmic ribosomes to stabilize ER translocation and ensure proper targeting of polypeptides destined for the ER lumen. TOM20 possesses a loop-like structure that resembles a similar portion of the TRAP receptor of the ER translocon, which is known to interact with the ribosomal protein rpL38A (Jaskolowski, Jomaa et al. 2023). Positioning an atomic model of the TOM complex with the TOM40 channel directly under the peptide exit tunnel and the globular domain of TOM20 at connection #2 positions this loop region near rpL38 at connection #3 (**Supplementary Figure 4D**), suggesting that perhaps similar interactions may occur to facilitate mitochondrial protein import.

In summary, our structure shows density for three contacts between the cytoplasmic ribosome and the OMM, which share structural similarities with ER-ribosome interactions during ER co-translational import. Identifying the similarities between these two systems expands our understanding of the potential molecular components mediating co-translational import into mitochondrial co-translational import.

### Cytoplasmic ribosomes primed for protein import cluster on the mitochondrial membrane surface

Previous cryo-ET analysis on purified mitochondria shows that mitochondrial-associated ribosomes cluster on the OMM (Gold, Chroscicki et al. 2017). We asked whether similar clustering is observed within the native cellular context. To quantify the degree of clustering, we used a recently developed software package called Tomospatstat (Martin-Solana, Diaz-Lopez et al. 2024) to analyze the three-dimensional spatial distribution of mitochondrial-associated ribosomes visible in our tomograms (**Figure 3, Supplementary Figure 5**). This program uses Ripley’s K function to measure how often different structures appear near each other and how close they are to their neighbors (Supplementary Figure 5A). For our analysis, the occurrence of the mitochondrial-associated cytoplasmic ribosomes on OMM in each tomogram within a given radius (r) is defined by K(r), and the occurrence that would be expected from complete spatial randomness (CSR) is defined by (KCSR(r)). In this analysis, the K(r)/KCSR(r) ratio represents the level of clustering within r, with values near 1 indicating that particles exhibit a random distribution across the space. We calculated and plotted curves for the K(r)/KCSR(r) ratio for mitochondrial-associated cytoplasmic ribosomes oriented for protein import within r values of 30-100 nm, which showed an overall trend towards clustering (curves higher than 1) (**Supplementary Figure 5C**). We plotted the maximum value of K(r)/KCSR(r) for given radius intervals of 10 nm for each tomogram. We found that mitochondrial-associated ribosomes oriented for import exhibit a higher degree of clustering on the OMM than ribosomes near but not oriented for protein import on the OMM (**Figure 3A, Supplementary Figure 5D**). The quantification was further confirmed visually by observing the clustering of co-translating ribosomes on the OMM at different K ratio values (**Supplementary Figure 5B**). This enhanced clustering was observed for radii from 30-90 nm, but most extreme at the closest radii interval of 30-40 nm (**Figure 3A**). This observation is consistent with previous reports of ribosomal clustering on purified mitochondria, which showed that 90% of ribosomes oriented for protein import form clusters less than 50 nm apart (Gold, Chroscicki et al. 2017).

**Figure 3.**
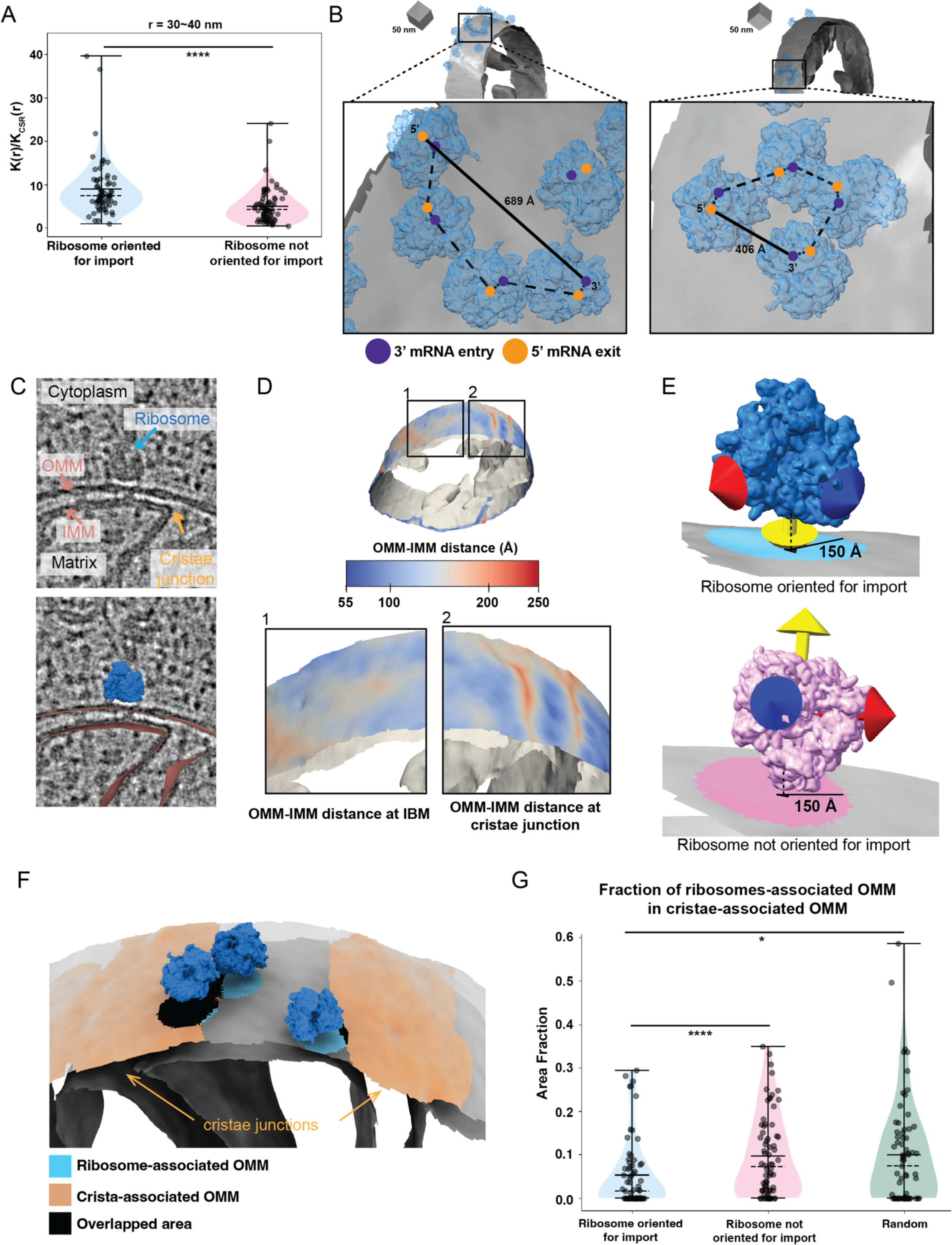
Cytoplasmic ribosomes primed for protein import cluster on the mitochondrial membrane. A. Quantification of the maximum value of K(r)/K_CSR_(r) for a 30-40 nm radius for each tomogram within the indicated ribosome class. P values from Mann-Whitney U test are indicated. *P < 0.05; **P < 0.01; ***P < 0.005; ****P < 0.001. B. Representative membrane surface reconstructions of mitochondria (gray) with ribosomes oriented for import relative to the OMM (blue). Insets show zoomed-in boxed regions of the ribosome models with circle overlays demarking the location of the 3’ mRNA entry (blue), the 5’ mRNA exit sites (orange), the possible pathways of interconnecting mRNA (dashed black line), and the calculated end-to-end distance from 5’ to 3’ of each interconnected mRNA (solid black line). C. Representative tomogram slices showing labeled cytoplasm, ribosome, mitochondrial matrix, IMM, OMM, and cristae junctions (upper panel) with an overlay of surface mesh reconstructions of IMM and OMM (red) and ribosome (blue) (lower panel). D. Representative membrane surface reconstruction of mitochondria with the OMM surface colored by outer-to-inner (OMM-IMM) membrane distance and the IMM surface shown in gray. The bottom inset labeled “1” shows inner membrane boundary (IBM) regions on OMM with more subtle OMM-IMM distance variations. In contrast, the bottom inset labeled “2” shows regions on OMM with large OMM-IMM distances corresponding to cristae junctions. E. Ribosome and membrane models defining the patches on the membrane surface mesh reconstruction that correspond to ribosomes oriented for import (blue, top) and ribosomes near but not oriented for import (pink, bottom). The ribosomes oriented for import are defined as those with the peptide exit tunnel (yellow arrow) pointed toward the membrane. In contrast, those not oriented for import have peptide exit tunnels facing away from the membrane. F. Representative ribosome and membrane model with the OMM surface colored by the ribosome-associated (blue) and crista-associated OMM (orange), with areas of overlap (black). G. Quantification of the average fraction of overlap from each tomogram between indicated ribosome class. P values from Mann-Whitney U test are indicated. *P < 0.05; **P < 0.01; ***P < 0.005; ****P < 0.001.

Clusters of ribosomes can form stable polysome structures that simultaneously translate the same mRNA transcript. These polysome structures have been observed on the ER membrane both *in vitro* and within the native cellular context to facilitate the co-translational import of polypeptides with signal recognition sequences into the ER lumen (Pfeffer, Brandt et al. 2012, Gemmer, Chaillet et al. 2023). We observe clusters of ribosomes oriented for import in formations that resemble polysome assemblies on the OMM with the mRNA entry and exit sites aligned across adjacent ribosomes (**Figure 3B, Supplementary Figure 6A**). By fitting rigid rods connecting the 3’ mRNA entry (blue circle in **Figure 3B, Supplementary Figure 6A**) and 5’ exit site (orange circle in **Figure 3B, Supplementary Figure 6A)** for each of the adjacent ribosomes, we can visualize the possible pathways of interconnecting mRNAs and calculate the end-to-end distance from 5’ to 3’ ends. The end-to-end distance of the putative mRNA of these polysomes ranges from ∼223-689 Å (**Figure 3B, Supplementary Figure 6A**), which falls in the range of the estimated 5’-to-3’ end-to-end length of mitochondrial-localized mRNAs associated with co-translational import (i.e., *TIM50*) (Guo, Modi et al. 2022). While this analysis suggests that these polysomes could accommodate a single mRNA transcript, future work is needed to determine whether clusters consist of single mRNAs with multiple ribosomes or independent mRNAs driving clustering.

Previous *in vitro* studies showed that ribosomes oriented for import are preferentially associated with regions of the OMM that overlap with crista junction portions of the IMM (Gold, Chroscicki et al. 2017). We, therefore, asked whether a similar spatial clustering was observed in the native cellular context. To automatically identify the regions of the OMM associated with crista junctions, we first used our Surface Morphometrics pipeline to calculate the distance between the IMM and OMM for each triangle within our surface mesh reconstructions (**Figure 3C & D**). We then visualized these distance measurements directly on the surface mesh reconstructions of the OMM of individual mitochondria within our tomograms (**Figure 3D)**. Consistent with our previous work (Barad, Medina et al. 2023), the resulting surface maps showed regions with the largest OMM-IMM distance were found at sites of crista junctions, enabling automated classification of these regions. We then identified the triangle on the OMM closest to the nearest triangle on the IMM that fell within the cristae junction region and calculated the “crista-associated” OMM patch as the triangles with a 150 Å radius (orange “patch” in **Figure 3F**). Next, we located the nearest triangle within the surface mesh reconstruction of the OMM to the subset of ribosomes oriented for import. We determined the triangles within a radius of 150 Å of this nearest triangle encompassed a “patch” of the membrane encompassing the entire footprint of cytoplasmic ribosomes engaged in import using visual inspection in ArtiaX (**Figure 3E**). We defined these regions as “co-translation ribosome-associated” patches (blue patch in **Figure 3E**). We repeated the same process for the ribosomes near but not oriented for import on the OMM (pink patch in **Figure 3E).** We calculated the fraction of overlap between the OMM patch closest to a ribosome (blue “patch” in **Figure 3F**) and the “crista-associated” OMM patch and plotted the average fraction of overlap (black “patch” in **Figure 3F)** from each tomogram as independent observations. We observed that the fraction of overlap of the cytoplasmic ribosomes optimally oriented for protein import is significantly lower relative to the degree of overlap for ribosomes near but not oriented for import on the OMM and random chance alone (**Figure 3G**).

This finding contrasts with previous reports, which showed that ribosomes oriented for protein import tend to cluster near cristae junctions on purified mitochondria (Gold, Chroscicki et al. 2017). One possible explanation for this discrepancy is that perhaps there are differences in the stability of ribosomes at different mitochondria regions. For example, ribosomes positioned near cristae junctions may be more tightly associated and, therefore, retained throughout the purification procedure. In contrast, those positioned in non-cristae junction regions may be less tightly associated and, therefore, de-stabilized during the purification procedure. Another possibility is that differences in mRNA localization on the mitochondrial surface may impact the stability and efficiency of co-translational import into the mitochondria. This may be further exacerbated by the transient cellular environment as a mechanism to regulate the import of specific proteins encoded by mRNAs at cristae junctions separately from the import at regions of the OMM corresponding to other IMM subdomains, such as the inner membrane boundary (IBM). Regardless, our study lays the groundwork for incorporating the contextual structural approaches described here to further decipher the molecular basis and regulation of mitochondrial protein import within the native cellular context.

Given the importance of inter-organellar contacts between the ER and mitochondria for various cellular functions (Csordás, Weaver et al. 2018, Koch, Lenhard et al. 2024), we wondered whether mitochondrial co-translational import might be preferentially associated with these regions. Interestingly, we only observed 1 example of an ER membrane that came within 25 nm of the OMM in our data, which exhibited no degree of overlap between the OMM associated with cytoplasmic ribosomes (**Supplementary Figure 6B**). Given the low occurrence of these contact sites within our data, this suggests that it is unlikely that ER-mitochondria contacts are regulating mitochondrial co-translational import. These analyses demonstrate that ribosomes primed for import tend to cluster on the OMM but not in regions associated with cristae junctions nor ER-mitochondria contact sites, suggesting that alternate mechanisms may dictate ribosomal localization on the mitochondrial surface in the native cellular context.

### Cytoplasmic ribosome-associated protein import alters the local architecture of the outer and inner mitochondrial membranes

Previous work demonstrated that the majority of co-translationally imported proteins are IMM that must be imported through translocases on both the OMM and IMM (Williams, Jan et al. 2014). We wondered whether the co-translational import is associated with local changes to mitochondrial membrane ultrastructure that may facilitate efficient protein translocation across these distinct mitochondrial compartments. Similar to the analysis used in **Figure 3E**, we used our Surface Morphometrics approach to define regions on the OMM as “co-translation-associated” patches (blue patch in **Figure 4A**). In contrast, all other triangles in the surface mesh reconstruction of the OMM were assigned as “non-co-translation-associated” patches (gray patch in **Figure 4A**). Next, we measured the distance to the closest triangle on the IMM mesh for each triangle within the OMM mesh for both “co-translation-associated” and “non-co-translation-associated” patches. We plotted a histogram of the combined distribution of distances for the surface mesh triangles within these patches, which showed an overall decrease in the intermembrane distance in the “co-translation-associated” patches relative to the “non-co-translation-associated” patches (**Supplementary Fig. 7A**). We plotted the mean OMM-IMM distance for each tomogram as independent observations and identified statistically significant decreases in the intermembrane spacing in “co-translation-associated” patches relative to “non-co-translation-associated” patches and relative to all membrane mesh triangles within each surface (**Figure 4B**). The quantification was further visually confirmed by observing that most co-translating ribosomes are found within OMM regions where the OMM-IMM distance is less than 120 Å (**Figure 4C**). As a negative control, we simulated random positions for the same number of “co-translation-associated” patches for each tomogram, which caused this significant decrease in local OMM-IMM distance to disappear (**Figure 4B)**. Despite notably fewer ribosomes optimally positioned for protein import in the vehicle-treated (i.e., non-CHX-treated), we observed a similar decrease in the intermembrane distance in “co-translation-associated” patches relative to “non-co-translation-associated” patches (**Figure 4B**).

**Figure 4.**
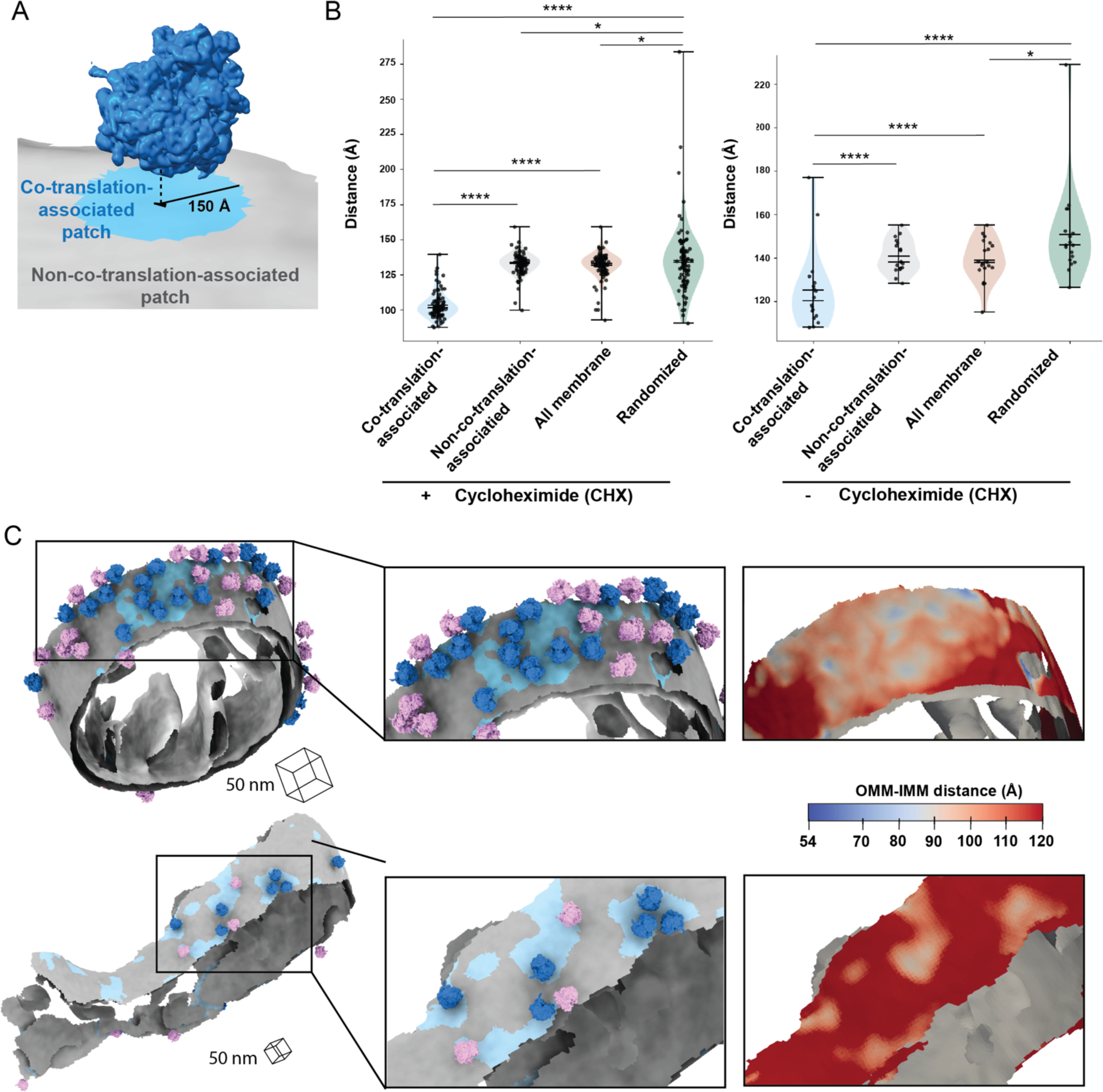
Ribosome-associated protein import alters the local architecture of the outer and inner mitochondrial membranes. A. Ribosome and membrane model defining co-translation-associated and non-co-translation-associated patches on membrane surface mesh reconstruction for OMM-IMM distance measurement. Co-translation-associated patches (blue) included the nearest OMM triangles (black in the middle of blue patch) to the import-oriented ribosomes and the OMM triangles within 150 Å of these nearest OMM triangles. Non-co-translation-associated patches (gray) consisted of the OMM mesh triangles that excluded co-translation-associated patches. B. Quantification of the peak histogram values of OMM-IMM distance measurements for each tomogram within the indicated membrane patch region. The co-translation-associated and non-co-translation-associated patches are detailed in Figure 4A. The “all membrane” patches represent the entire OMM surface. The “randomized” patches were simulated according to the number of co-translation-associated patches, as outlined in ***Materials and Methods***. P values from Mann-Whitney U test are indicated. *P < 0.05; **P < 0.01; ***P < 0.005; ****P < 0.001. C. Representative membrane surface reconstruction of mitochondria colored by OMM-IMM membrane distance, with regions less than 10 nm shown in blue and regions greater than 10 nm shown in gray. Ribosomes oriented for import relative to the OMM are colored blue, and the remaining ribosomes near but not oriented for import are shown in pink. Insets show zoomed-in boxed regions of the models (middle) and the local variations in OMM-IMM distance (right).

Overall, these contextual morphometrics analyses show decreases in the local intermembrane spacing at “co-translation-associated” membrane regions in both the presence and absence of CHX treatment. This suggests that local membrane remodeling may facilitate efficient protein import and translocation of nuclear-encoded mitochondrial proteins across the OMM and IMM. Notably, these local decreases in OMM-IMM distances at ribosome-associated import sites align with previous reports of intermembrane spacing detected using quantum dots to locate protein import mediated by the TOM-TIM23 complex (Gold, Ieva et al. 2014). The similarity in the distance measurements between the previous work and this current study suggests that the OMM-IMM distance required for protein import likely falls within this range. In previous cryo-ET investigations of ribosomal-OMM interactions on purified mitochondria, the OMM-IMM distance in non-protein importing areas was found to be around the same range as the protein importing areas (Gold, Ieva et al. 2014), indicating no membrane remodeling for priming protein import. In contrast, we observed that the average distance between the OMM and IMM in areas not associated with ribosomes oriented for protein import is around 130-140 Å, which is greater than the measured OMM-IMM distance observed *in vitro.* This discrepancy could be due to non-native membrane rearrangements that occur during the purification process that are absent in the native cellular context or limitations in the accuracy of the quantification procedures used previously relative to the Surface Morphometrics approach used in this current study. Further, this highlights the importance of probing these functional interactions in the native cellular context.

## CONCLUSIONS

Recent advancements in cellular cryo-electron tomography imaging have enabled new avenues to structurally investigate cellular proteomes functioning in their native environment (Young and Villa 2023). This work takes advantage of several of these new developments across multiple steps in the cellular cryo-electron tomography workflow to structurally characterize the molecular architecture of cytoplasmic ribosomes at the surface of mitochondria (**Figure 1)**. We show here that the surface mesh reconstructions generated using our Surface Morphometrics approach (Barad, Medina et al. 2023) can be used to identify a subset of macromolecules positioned in orientations relative to cellular membranes that are crucial for performing their biological function (**Figure 1F-I, Supplementary Figure 2**). We used this approach to identify, align, and average a subset of cytoplasmic ribosomes optimally oriented and positioned for protein import (**Figure 2, Supplementary Figures 3 & 4**). To our knowledge, we present the first subtomogram average of a cytoplasmic ribosome forming three distinct connections with the OMM surrounding the peptide exit tunnel (**Figure 2).** We further harnessed advanced morphometrics approaches to quantify local changes to membrane ultrastructure at regions associated with import, including changes to OMM-IMM distance and the degree of clustering on the mitochondrial surface (**Figure 3 & 4, Supplementary Figure 5-7**). Overall, this work sets the stage for enabling exciting opportunities to identify the molecular players regulating these interactions and local remodeling during mitochondrial protein import using genetic knockdown approaches.

## MATERIALS AND METHODS

### Yeast strains and growth conditions

The yeast strain used in this study is a derivative of the *Saccharomyces cerevisiae* strain BY4741 which contains Su9-mCherry-Ura3 and TIM50-GFP-His3MX6. They were recovered by streaking on the fresh YPD agar (1% yeast extract, 2% peptone, 2% glucose, 2% agar) plates and incubated at 30°C for ∼2 days. Yeast single colony on the plate was inoculated in YPD (1% yeast extract, 2% peptone, 2% glucose) or YPG (1% yeast extract, 2% peptone, 2% glycerol) and grown overnight at 30°C. The overnight culture was diluted to OD600 of 0.2 with the corresponding medium and then grown to OD600 of 0.8 at 30°C before vitrification.

### Sample preparation for cryo-ET

Yeast liquid cultures with OD600 of 0.8 was 4 times diluted to OD600 of 0.2 with the medium supplemented with 133 μg/mL cycloheximide (CHX). The final concentration of CHX was 100 μg/mL. Yeast liquid cultures were incubated with CHX for 2 minutes, and then 4 μL of the sample was applied to the glow-discharged R1/4 Carbon 200-mesh gold EM grid (Quantifoil Micro Tools). The EM grid was incubated in the chamber of Vitrobot (Vitrobot Mark 4; Thermo Fisher Scientific) for another 2 minutes before it was plunge-frozen in a liquid ethane/propane mixture. The Vitrobot was set at 30°C with 100% humidity, and the blotting was performed manually from the back side of grids using Whatman #1 filter paper strips through the Vitrobot chamber side port.

### Cryo-FM for examining sample quality

Vitrified grids were clipped in Cryo-FIB Autogrid (Thermo Fisher Scientific) in the vapor of liquid nitrogen (LN2). The clipped grids were loaded into the stage of Leica CryoCLEM microscope (Leica) to acquire the fluorescence/bright-field tiled image maps (atlases) in cryo condition by Leica LAS X software (25 µm Z stacks with system optimized steps, GFP channel ex: 470, em: 525). Z stacks were stitched together as a maximum projection map by LAS X navigator for examining the cell density and ice quality on EM grids.

### Cryo-FIB-milling for lamella generation

Cryo-FIB-milling of lamella was performed using Aquilos 2 cryo-FIB/SEM (Thermo Fisher Scientific) operated by software xT (Thermo Fisher Scientific). The atlases from cryo-FM were loaded into MAPS software (Thermo Fisher Scientific) to overlay with the SEM atlases of the same grids. Before milling, EM grids were first subjected to a layer of platinum sputter for 15 s (1 kV, 20 mA, 10 Pa). Next, the grids were coated with an organometallic platinum layer using a gas injection system for 45 s, and finally sputter-coated for 15 s (1 kV, 20 mA, 10 Pa). The regions of interest were selected in MAPS and then transferred to AutoTEM for identification of eucentric position, beam shifts, and tilt values in the preparation step. Before automated milling, the exact lamella positions were defined by the milling pattern with the width and height of 10 µm and thickness of 250nm. After assigning the position of the lamella, the relief cuts were generated by a relatively high ion current (0.5 nA) with a width of 1 µm and a height of 6.5 µm. Next, the automated milling in AutoTEM was executed to remove bulk cellular material. The automated milling task was separated into three different steps. In the rough milling step, 0.5 nA ion current was applied to generate 2 µm thickness of lamella with 13 µm front width and 12 µm rear width. In medium milling step, the 0.1 nA ion currents was applied to generate 1.2 µm thickness of lamella with 11.3 µm front width and 11 µm rear width. In the fine milling step, 50 pA was applied to generate 600 nm thickness of lamella with 10.1 µm width. After the milling steps, the thinning step was automatically executed with an ion beam of 50pA to generate 300 nm thickness of lamella with 10 µm width. A second thinning step was used to generate the 250 nm thickness of lamella with 10 µm width using 50 pA ion beam. After the automatic milling and thinning process, a polishing step was manually executed using an ion beam of 50 pA and targeted for the thickness of lamella under 200 nm.

### Tilt series collection

Tilt series collection on the lamella was performed on Titan Krios (Thermo Fisher Scientific) operated at 300 keV and equipped with a K3 direct electron camera and a BioQuantum energy filter (Gatan). Individual lamellae were montaged with low dose (1 e/Å^2^) at high magnification to localize cellular features and identify mitochondria by their distinctive OMM and IMM. Target selection and data acquisition were performed by Parallel cryo-electron tomography (PACE-tomo) (Eisenstein, Yanagisawa et al. 2023), which is a set of Python-based SerialEM scripts allowing multiple tilt series collection in parallel on the same lamellae via beam shift. Data was acquired at a magnification of 53,000x with a pixel size of 1.6626 Å or 33,000x with a pixel size of 2.638 Å and a nominal defocus range between -4 to -6 µm with 1 µm step. Tilt series collection was done in a dose-symmetric scheme with 3° tilt increment and angles ranging from -60° and +60° centered on -11° pretilt. Data was collected with dose fractionation, with 10∼11 0.28∼0.33 e/Å^2^ frames collected per second. The total dose per tilt is ∼3 e/Å^2^ and the total accumulated dose for the tilt series was ∼120 e/Å^2^.

### Tilt series processing and reconstruction

Tilt series micrograph movies were pre-processed through contrast transfer function (CTF) estimation and motion correction in Warp (Tegunov and Cramer 2019). Next, the pretilt angle applied during data collection was adjusted to 0° by a homemade script (adjust_mdoc.py) before the pre-processed tilt series output as averaged stacks in Warp. The averaged stacks were first aligned by batchtomo scripts in etomo using patch-tracking with four times binning (Mastronarde 1997). Next, the contour of alignment was manually curated by removing the poorly aligned patches. The aligned stacks were then imported into Warp for tomogram reconstruction with six times binning. This resulted in a voxel size of (9.98 Å)^3^ and volumes of 682 × 960 × 334.

### Template matching for localizing ribosome particles in tomograms

Cytoplasmic ribosome particles were localized by template-matching reconstructed tomograms against a yeast 80S ribosome (EMD-11096) by PyTom (Maurer, Siggel et al. 2024). The template was modulated with the CTF function parameters according to the defocus of each tomogram at 300 keV voltage, 2.7 mm Cs, and 0.1 amplitude contrast. Then the templates were low-pass filtered to 25 Å followed by downsampling to the pixel size of 9.98 Å. The final box size of templates was 45^3^ voxel size. All template modulation was performed by “pytom_create_template.py” script. A spherical mask with a radius of 22-pixel size with a soft edge was created later by the “pytom_create_mask.py” script. Template matching was performed using an angular search of 7° using the “pytom_match_template.py” script. Low-pass and high-pass filters of 25 Å and 500 Å were respectively applied to templates and tomograms. Initial particle candidates were extracted by “pytom_extract_candidates.py” script with the maximum number of particles of 2000. The optimized parameters with a 22-voxel masking radius and 10 number-of-false-positive were applied to perform the final extraction of ribosome particles as .star files.

### Subtomogram averaging analysis for cytoplasmic ribosomes

The annotation of cytoplasmic ribosome particles (.star files) was imported into Warp for subtomogram extraction. The subtomograms were initially extracted using 3x binning, resulting in a voxel size of (4.99 Å)^3^. The subtomograms, CTF volumes, and the star file were transferred to Relion (4.0b1-cuda) for subtomogram averaging analysis (Zivanov, Otón et al. 2022). The initial alignment and averaging were performed using Relion reconstruction (relion_reconstruct_mpi). This initial model was further used as a reference for alignment and refinement in Relion auto-refine (relion_refine_mpi). The aligned particles in Relion were further refined in M using the two half maps as references (Tegunov, Xue et al. 2021), resulting in a resolution of 10.3 Å (∼Nyquist). The subtomograms aligned with M were extracted using 2x binning, resulting in a voxel size of (3.33 Å)^3^. The 2x binning subtomograms from M were processed by the same Relion-M workflow as above. The final resolution of the cytoplasmic ribosome was determined to be 8 Å by the Fourier Shell Correlation (FSC) score of 0.143.

### Membrane tracing, voxel segmentation, and surface generation

The reconstructed tomograms with six times binning (voxel size: (9.98 Å)^3^) were processed by Membrain-Seg (Lamm, Zufferey et al. 2024), which is an advanced machine-learning software based on UNet for tracing and segmenting cellular membranes. The traced volumes of membranes were imported into AMIRA (Thermo Fisher Scientific) for manual curation. OMM and IMM were designated as different labels by 3D Magic Wand, and the ambiguous connection between them was manually curated slice-by-slice using the 2D Brush tool. Next, the individual labels of membrane voxel segmentation were reconstructed as smooth surfaces using the “segmentation_to_meshes.py” script with Angstrom units in Surface Morphometrics (Barad, Medina et al. 2023). The surfaces were generated with a maximum of 150,000 triangles, a reconstruction depth of 8, an interpolation weight of 8, and an extrapolation distance of 25 Å. The surface orientations were further refined using the “run_pycurv.py” script in surface morphometrics.

### Calculation of the relative distance and orientation between ribosomes and OMM surfaces and the relative distance between ribosome peptide tunnel exits and OMM surfaces

We used the relative distance and the angle to represent the relative localization and orientation between ribosomes and the nearest surface triangles. In the final refinement run, the starfile provided the coordinates (rlnCoordinateX, rlnCoordinateY, and rlnCoordinateZ) and orientation as Euler angles (rlnAngleRot, rlnAngleTilt, and rlnAnglePsi) of cytoplasmic ribosomes. Additionally, the Surface Morphometrics pipeline provided the coordinates and orientation as normal vectors of surface triangles in triangle graph files (.gt). To determine the relative distance between ribosomes and OMM surfaces, we incorporated the coordinates from these data and utilized a k-dimensional tree Python function to calculate the minimum distance from each ribosome to the nearest OMM surface triangle. To calculate the relative angle between ribosomes and the nearest OMM surface triangles, we first converted the Euler angles of ribosomes to rotation matrixes through a Python function “eular2matrix” and then extracted the vectors which visually represented as the three-color arrows in ArtiaX (Ermel, Arghittu et al. 2022), which is a plugin in ChimeraX (Pettersen, Goddard et al. 2021). The relative angle between each vector of the ribosome and the normal vector of the nearest OMM surface triangle was calculated as the following equation:

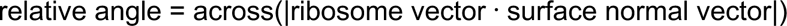

We positioned a spherical mask that covers the peptide exit of the 8 Å yeast 80S ribosome structure in ChimeraX and then determined the center of the mask in 3dmod (Kremer, Mastronarde et al. 1996). We then shifted the center of ribosome to the center of the peptide exit for all ribosome particles and extracted the particles to get the starfile with shifted coordinates using M. The relative distance between the peptide exit of ribosomes and the nearest OMM surface triangles was determined by the k-dimensional tree in Python as well.

The relative distance between ribosomes and OMM surfaces, the three relative angles between ribosomes and OMM surfaces, the identifiers of the nearest OMM surface triangles, and the relative distance between the peptide exit of ribosomes and OMM surfaces were recorded in the CSV files per tomogram for further analysis. This calculation is accomplished by “ribo_membrane_distance_orientation.py” script. The CSV files were further converted to starfiles per tomogram by the “csv_to_star.py” script for rendering particles in ArtiaX. The optimal cutoff for identifying the ribosomes that oriented for protein import on mitochondria was determined manually using ArtiaX, with a final threshold of 95 Å for the distance between peptide exit and OMM.

### Subtomogram averaging analysis for cytoplasmic ribosomes optimally oriented for protein import on mitochondrial surfaces

The starfile was filtered based on a final threshold of 95 Å for the distance between the peptide exit and OMM to select the ribosomes optimally oriented for protein import into mitochondria. The orientation of selected ribosomes was manually examined in ArtiaX. This filtered and curated starfile was then input into Relion (4.0b1-cuda) for initial alignment and averaging using Relion reconstruction (relion_reconstruct_mpi). The initial model was subsequently used as a reference for alignment and refinement in Relion auto-refine (relion_refine_mpi), resulting in a resolution of 19 Å. Various low-pass filters were applied to the output model from Relion auto-refine using relion_image_handler to visualize the connection between the cytoplasmic ribosome and the mitochondrial membrane (***Supplementary Figure 3A***). The S. cerevisiae 80S ribosome (PDB 4V6I) was fitted into the 30 Å low-pass filtered map to study the ribosomal components within the map. Finally, the model from Relion auto-refine was further postprocessed in M to estimate the final resolution, determined by a Fourier Shell Correlation (FSC) score of 0.143.

### Subtomogram averaging analysis for cytoplasmic ribosomes that are near but are not optimally for protein import on mitochondria

The starfile was filtered based on the following criteria to select the ribosomes that are in proximity to mitochondria but are not oriented for protein import: (1) the distance between ribosomes and OMM is less and equal to 250 Å (2) the coordinates are not included in the population of ribosomes that oriented for import. The filtered starfile was input into Relion for reconstruction (relion_reconstruct_mpi). The reconstructed model was used as reference in Relion auro-refine (relion_refine_mpi).

### Analysis of spatial clustering of mitochondria-associated cytoplasmic ribosomes

We examined the clustering patterns of mitochondria-associated cytoplasmic ribosomes using Tomospatstat (Martin-Solana, Diaz-Lopez et al. 2024), which employs Ripley’s K function K(r) to describe the occurrences of objects within certain distances r. In this analysis, the occurrence K(r) of mitochondrial-associated cytoplasmic ribosomes on the OMM in each tomogram was calculated and compared with the function K of complete spatial randomness (KCSR(r)). The K(r)/KCSR(r) ratio indicates the level of clustering within r. We calculated this ratio for mitochondrial-associated cytoplasmic ribosomes oriented for protein import and for cytoplasmic ribosomes 250 Å away from mitochondria but not oriented for protein import, within r values ranging from 27-166 nm. The parameters we used in Tomospatstat were as follow:

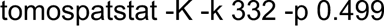

The K(r)/KCSR(r) ratios along r for these two ribosome populations were recorded as CSV files for each tomogram. The curves of K(r)/KCSR(r) ratios along r were plotted using Matplotlib. The maximum values of K(r)/KCSR(r) for given radius intervals of 10 nm for each tomogram were visualized using violin plots. The Mann– Whitney U test was applied to assess the statistical significance of differences in the maximum values of K(r)/KCSR(r) between the two ribosome populations. The generation of violin plots and statistical test were performed by “plotting_tomospatstat_K_ratio_stats.py “.

### Identifying polysomes on mitochondrial membranes

We identified the positions of the 5’ mRNA exit and 3’ mRNA entry in our 8 Å cytoplasmic ribosome map by fitting the model of S. cerevisiae 80S ribosome (PDB 4V6I) in ChimeraX. Using the “Map Eraser” tool in ChimeraX, we cropped a sphere with a 20 Å radius from the map of the cytoplasmic ribosome at the 5’ mRNA and 3’ mRNA positions, saving these sphere maps to mark the locations of the 5’ mRNA exit and 3’ mRNA entry.

To display the optimally oriented cytoplasmic ribosomes along with their 5’ mRNA exit and 3’ mRNA entry on the mitochondrial surface, we loaded the surface files (.stl) of the OMM and IMM, and the starfile of optimally oriented cytoplasmic ribosomes rendered with the maps of the cytoplasmic ribosome, 5’ mRNA exit, and 3’ mRNA entry in ArtiaX. We then manually searched for potential polysomes using the following criteria: (1) the nearest neighboring ribosome is within 30 nm, and (2) the 3’ mRNA entry is directly adjacent to the 5’ mRNA exit of the nearest neighboring ribosome. To show the interconnecting pathway of mRNA associated with polysomes, we placed markers at the 5’ mRNA exit and 3’ mRNA entry for each ribosome and connected the markers with dashed rods in the order of mRNA from 5’ to 3’ using the distance measurement tool in ChimeraX. We measured the end-to-end distance from 5’ to 3’ of these polysome-associated mRNAs using the distance measurement tool in ChimeraX.

### Analysis of the overlap fraction of “ribosomes-associated OMM” in “crista-associated OMM”

We identified the “ribosome-associated OMM” regions for cytoplasmic ribosomes oriented for protein import and for those 250 Å away from the OMM but not oriented for protein import. To pinpoint these OMM regions, we first extracted the identifiers of the nearest OMM surface triangles for these ribosomes using the “match_particles.py” script. We then used these identifiers to locate the coordinates of the nearest OMM surface triangles from the OMM triangle graph files (.gt) and searched for triangles within a radius of 150 Å around these nearest OMM surface triangles using “OMM-patches_IMM_dist_measurement.py”. We saved two separate OMM triangle graph files (.gt) with the label as “ribosome-associated OMM” for these two populations of ribosomes. Randomized ribosome-associated OMM regions was generated for each tomogram by “random_patches_OMM-IMM_dist_measurement.py” with the criteria as follows: (1) the number of randomized OMM regions matched the number of “ribosome-associated” OMM regions that associated with the cytoplasmic ribosomes oriented for import, and (2) the distances between the centers of the randomized regions were greater than 150 Å. The separate OMM triangle graph files (.gt) with the label as “ribosome-associated OMM” were also saved for randomization.

To identify crista-associated OMM, we first subclassified cristae junctions by measuring the distance from the IMM to the OMM using Surface Morphometrics with the “measure_distances_orientations.py” script. We selected the IMM triangles that are 18-30 nm away from the OMM and manually cleaned up the selected IMM triangles that did not belong to cristae junctions. We extracted the subclassified IMM with manual curation as CSV files, which contained the identifiers of the nearest OMM surface triangles in Paraview. We then added a new label as cristae junction projected OMM for the OMM triangles with these identifiers in OMM triangle graph files (.gt) using “CJ_projected_OMM.py”. Next, we searched the OMM triangles within 15 nm (half the width of a cristae body) of the cristae junction projected OMM and added a new label as “crista-associated OMM” for these OMM triangles in the OMM triangle graph files (.gt) using “expand_CJ_projected_OMM.py”.

To quantify the overlap fraction of ribosome-associated OMM in crista-associated OMM per tomogram, we first counted the area of the OMM triangles with the label of “ribosome-associated OMM”, and the area of the OMM triangles with both labels of “ribosome-associated OMM” and “crista-associated OMM” from OMM triangle graph files (.gt). We then calculated the overlap fraction as follows by “count_overlap_area.py”: Overlap fraction = (Area of the OMM labeled with “ribosome-associated OMM” and “crista-associated OMM”) / (Area of the OMM labeled with “ribosome-associated OMM”)

The overlap fraction per tomogram for both populations of ribosomes and the randomization was plotted as violin plot. The Mann–Whitney U test was applied to assess the statistical significance of differences in the overlap fraction. The generation of violin plot and statistic test were performed using “plotting_overlap_fraction_stats.py”.

### Calculation of distances between OMM and IMM at “co-translation-associated” and “non-co-translation-associated” patches on the OMM

We defined “co-translation-associated” patches as areas where ribosomes oriented for import are associated with the OMM surface. To identify these patches, we first extracted the identifiers of the nearest OMM surface triangles for these ribosomes using the “match_particles.py” script from the CSV files by putting the starfile. We then used these identifiers to locate the coordinates of the nearest OMM surface triangles from the OMM triangle graph files (.gt) and searched for triangles within a radius of 150 Å around these nearest OMM surface triangles as ribosome-associated patches. The distances between ribosome-associated patches and IMM were further calculated using the k-dimensional tree in Python. The identification of “co-translation-associated” patches and the distance calculation were accomplished by “OMM-patches_IMM_dist_measurement.py”. On the other hand, “outside_OMM-patches_IMM_dist_measurement.py” was applied to extract “non-co-translation-associated” patches by excluding “co-translation-associated” patches, and to calculate the distances between “non-co-translation-associated” patches on the OMM and IMM. Randomized co-translation-associated patches were generated for each tomogram by “random_patches_OMM-IMM_dist_measurement.py” based on the following criteria: (1) the number of randomized patches matched the number of “co-translation-associated” patches, and (2) the distances between the centers of the randomized patches were greater than 150 Å. The distance calculation between the complete OMM and IMM was performed by Surface morphometrics using “measure_distances_orientations.py”. Violin plots were created by generating histograms with 100 bins for each tomogram and identifying the peak value of the most populated bins. The Mann–Whitney U test was used to analyze the statistically significant differences in peak positions. The generation of violin plots and the statistical test were accomplished by “plotting_OMM-patches_IMM_dist_stat.py”.

## Data and Code Availability

All tilt series, reconstructed tomograms, voxel segmentations, and reconstructed mesh surfaces used for quantifications were deposited in the Electron Microscopy Public Image Archive (EMPIAR) under accession codes EMPIAR-XXXXX. All subtomogram averages were deposited in the Electron Microscopy Data Bank (EMDB) under accession codes EMDB-XXXX. All scripts used for Surface Morphometrics are available at https://github.com/grotjahnlab/surface_morphometrics).

## ACKNOWLEDGEMENTS

We thank Bill Anderson and William Lessin at The Scripps Research Institute Hazen cryo-electron microscopy facility for microscope support and Jean-Christophe Ducom at The Scripps Research Institute for computational support. We also thank R. Luke Wiseman for their critical input on the manuscript. D.A.G. is supported by Nadia’s Gift Foundation Innovator Award of the Damon Runyon Cancer Foundation (DRR-65-21) and the National Institutes of Health (NIH) grant RF1NS125674. B.M.Z is supported by the National Institutes of Health (NIH) grant R35GM128798. This work used equipment supported by NIH grant S10OD032467.

## AUTHOR CONTRIBUTIONS

**Y. Chang:** Conceptualization, Data curation, Formal analysis, Investigation, Methodology, Software, Validation, Visualization, Writing - original draft, Writing - reviewing and editing. **B.A. Barad:** Methodology, Software, Writing - reviewing and editing. **H. Rahmani:** Methodology, Software, Writing - reviewing and editing. **B.M. Zid:** Conceptualization, Funding acquisition, Resources, Writing - reviewing and editing. **D. A. Grotjahn:** Conceptualization, Funding acquisition, Methodology, Project administration, Resources, Supervision, Validation, Visualization, Writing - original draft, Writing - reviewing and editing.

## SUPPLEMENTARY FIGURES

**Supplementary Figure 1.**
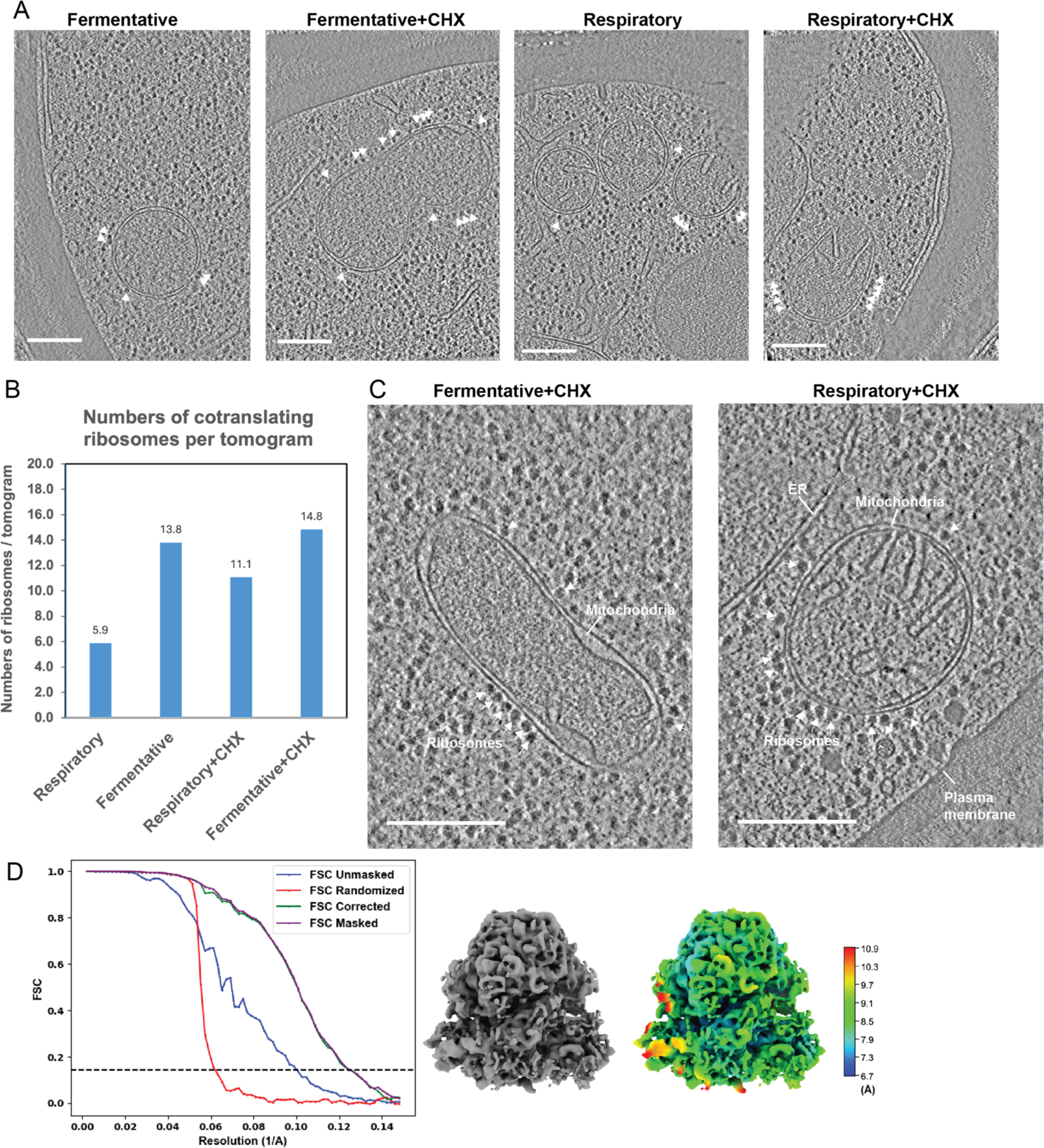
Representative tomograms of cryo-focused ion beam (cryo-FIB) tomograms milled *S. cerevisiae* cell lamellae with visible mitochondria-associated cytoplasmic ribosomes. A. Representative X-Y slices of reconstructed tomograms collected at pixel size 2.638 Å from cryo-FIB milled *S. cerevisiae* yeast cells grown in different growth conditions (i.e., fermentative and respiratory) and treatment conditions (i.e., vehicle and cycloheximide, CHX). Cytoplasmic ribosomes in close proximity to the OMM are highlighted by white arrowheads. Scale bars = 250nm B. Quantification of the number of ribosomes positioned with the exit tunnel facing the OMM in CHX-treated or vehicle-treated cells grown in respiratory versus fermentative conditions. C. Representative X-Y slices of reconstructed tomograms collected at pixel size 1.6626 Å from cryo-FIB milled *S. cerevisiae* grown in fermentative and respiratory conditions and treated with CHX (100 μg/mL) displaying subcellular features such as mitochondria, ribosomes, the endoplasmic reticulum, and the plasma membrane. Scale bars = 250 nm D. Fourier shell correlation plot (left) of the 80S cytoplasmic ribosome reconstruction is shown with resolution reported at 0.143 FSC and the reconstructed subtomogram average (middle) shown to the right of the curves. The 80S cytoplasmic ribosome was resolved to 8 Å from 35,784 ribosome particles with the color map (right) shows the local resolution.

**Supplementary Figure 2.**
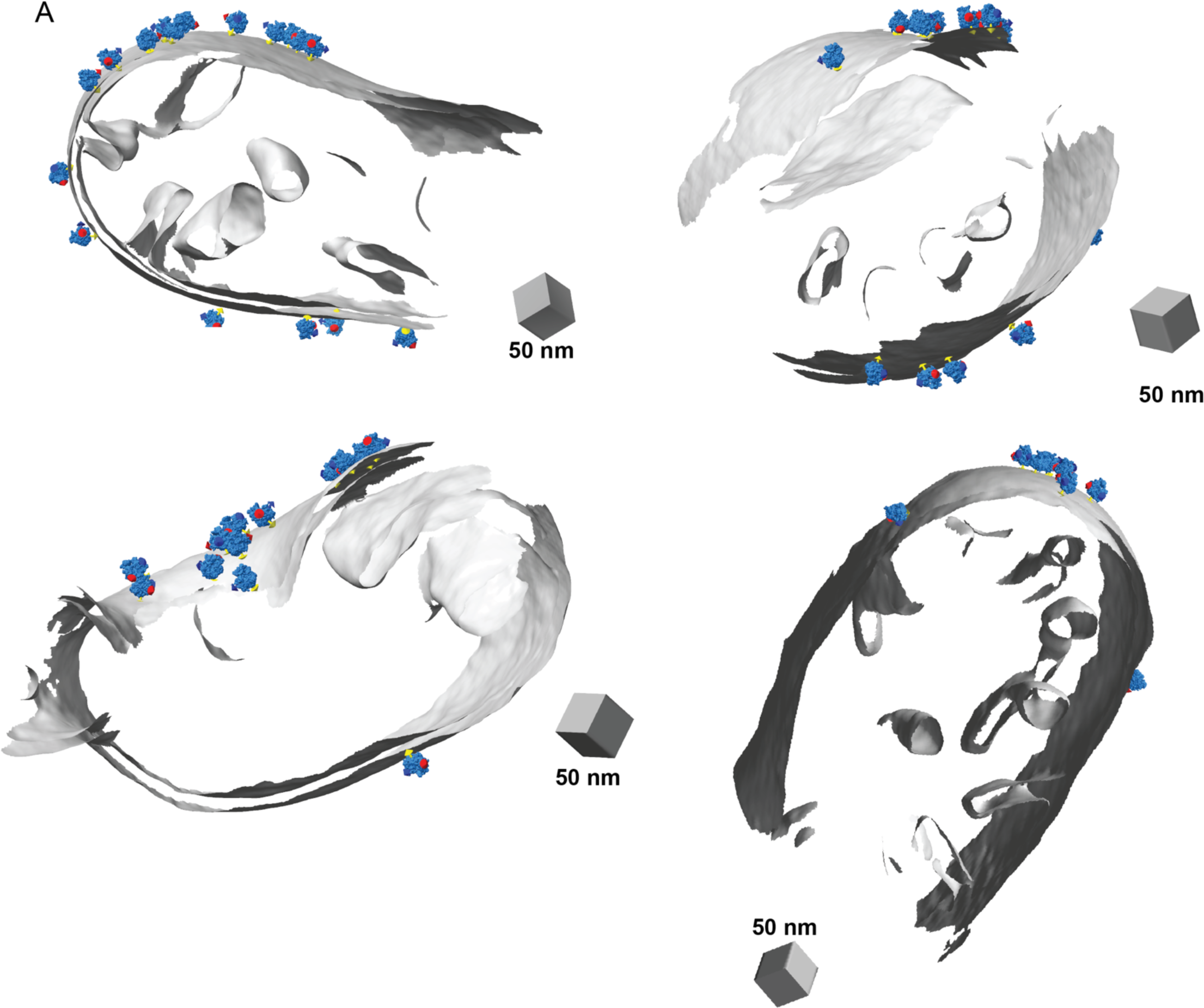
Distance filtering identifies a subset of cytoplasmic ribosomes optimally positioned for protein import into the OMM. A. A subset of representative models of ribosomes positioned with their exit tunnels optimally positioned for protein import on the OMM surface.

**Supplementary Figure 3.**
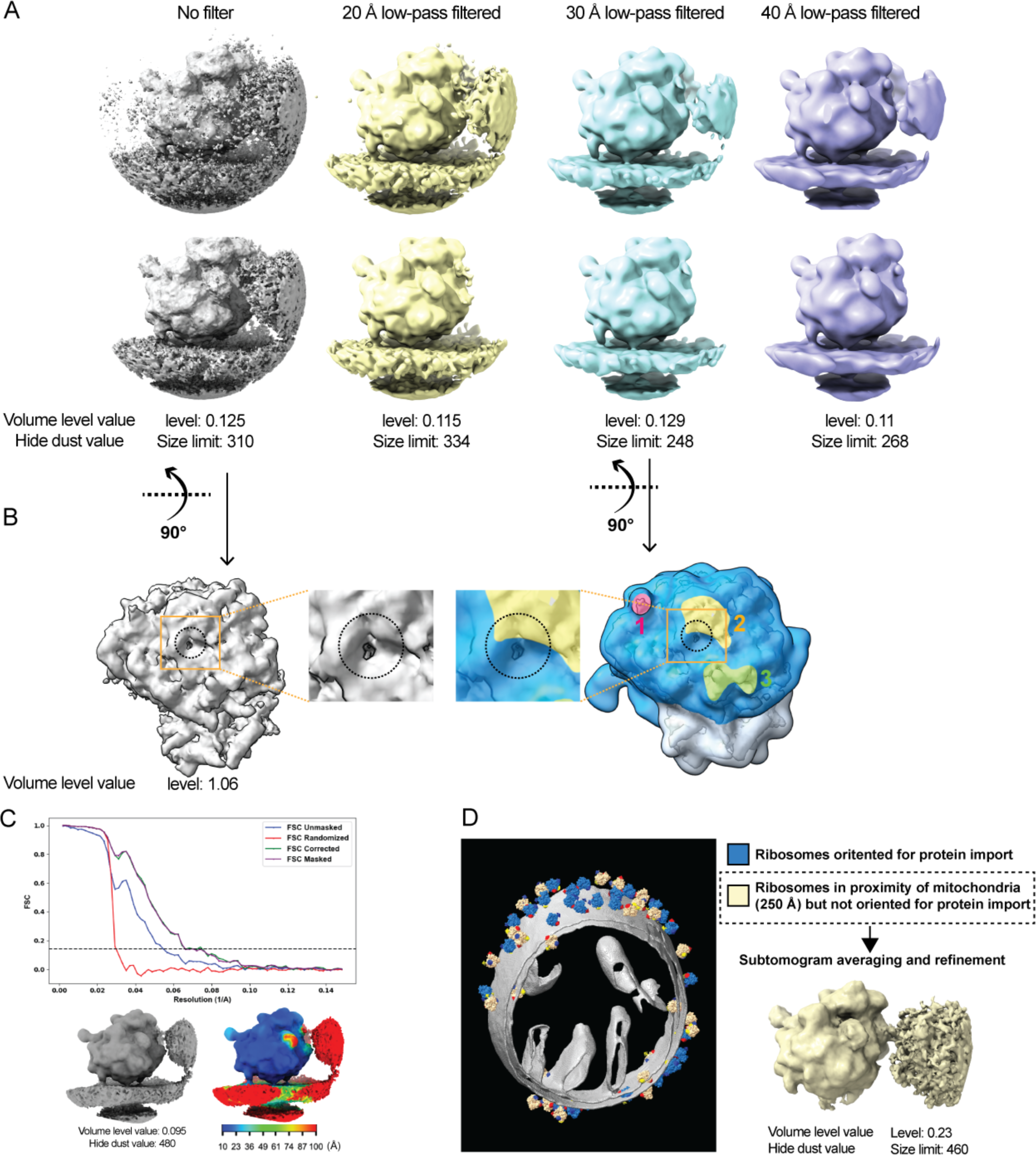
Three-dimensional subtomogram average of a cytoplasmic ribosome positioned for protein import on the outer mitochondrial membrane (OMM). A. Subtomogram average of a cytoplasmic ribosome positioned for protein import on the OMM displayed with varying levels of low-pass filter, isosurface volume threshold, and hide dust values in ChimeraX. The densities surrounding the ribosome on the cytoplasmic side, as observed in other studies (Brandt, Carlson et al. 2010, Pfeffer, Brandt et al. 2012, Gemmer, Chaillet et al. 2023), likely correspond to neighboring ribosomes. B. The peptide exit tunnel is visible in the subtomogram average (gray density) at lower isosurface volume thresholds, as indicated by the dashed black line. This was used to mark its position relative to the connecting densities visible in the subtomogram average at higher isosurface volume threshold values (colored density). C. Fourier shell correlation plot of the OMM-associated 80S cytoplasmic ribosome reconstruction is shown with resolution reported at 0.143 FSC and the reconstructed subtomogram average (bottom left) shown to the right of the curves. The 80S cytoplasmic ribosome was resolved to 19 Å from 1,076 ribosome particles with the color map shows the local resolution (bottom right). D. Representative models of ribosomes positioned within 250 Å of the OMM surface with their exit tunnels facing away from the OMM. 3D refinement of these particles results in 3D reconstructions of a ribosome that does not contain any distinguishable connecting densities between the 80S ribosome and the OMM, suggesting that these connections are specific to 80S ribosomes optimally positioned for protein import.

**Supplementary Figure 4.**
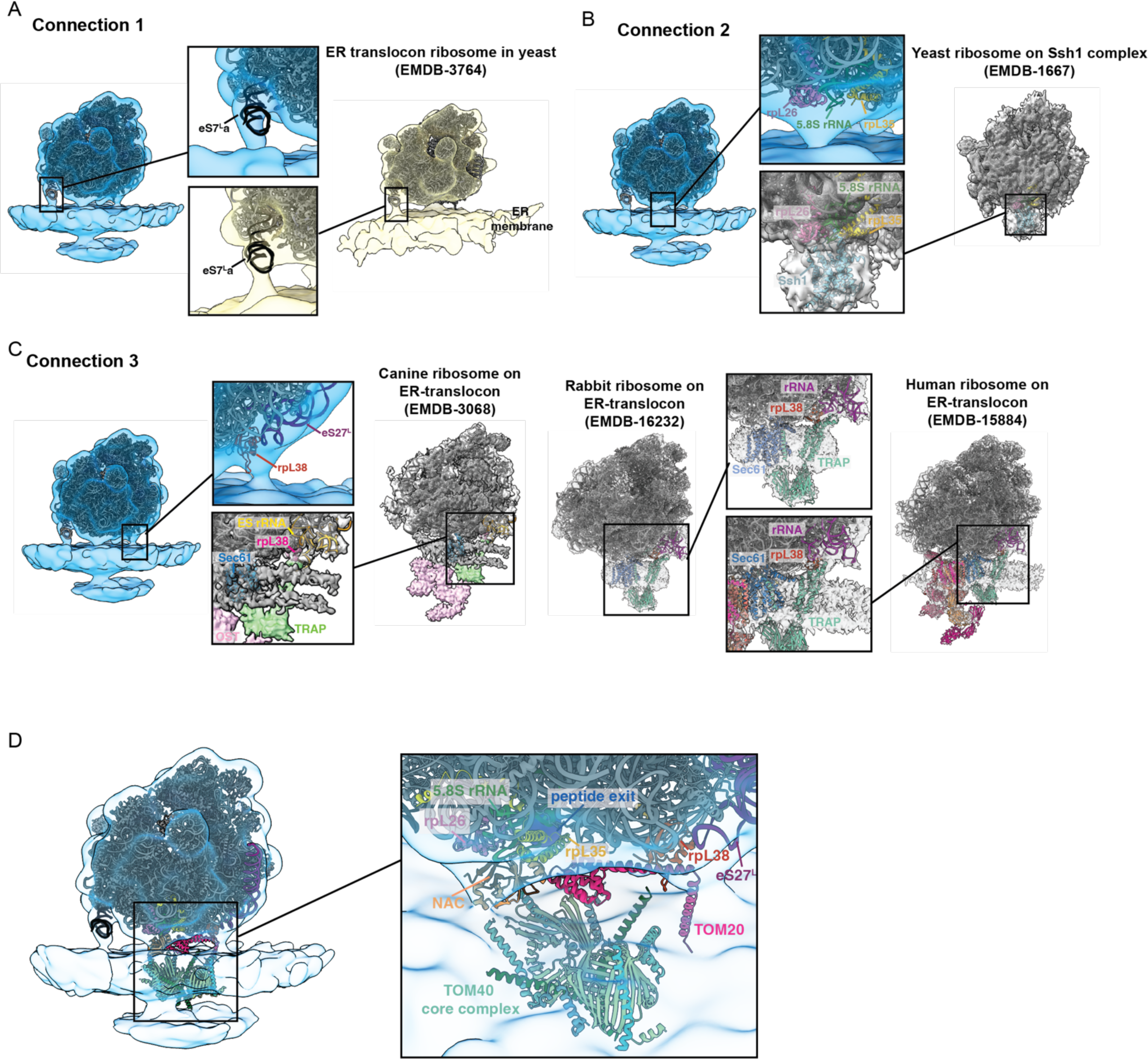
Similarities between the organization of the mitochondrial-associated ribosome and ER-associated ribosome structures, and the hypothesized model for the architecture of mitochondrial co-translational import. A. Density corresponding to the connection labeled #1 in the subtomogram average of mitochondrial-associated ribosomes correlates well with the density corresponding to the expansion segment of eS7La of the 25S rRNA in the large 60S subunit present in the ER-associated ribosome maps from *S. cerevisiae* (EMD-3764). B. Density corresponding to the connection labeled #2 correlates well with the density corresponding to the region where ribosome interacts with the import channel, Ssh1, in the ER-associated ribosome maps from *S. cerevisiae* (EMD-1667). C. Density corresponding to the connection labeled #3 correlates well with the density corresponding to the rRNA expansion segment, rpL28, and the translocon-associated protein complex (TRAP) in the ER-associated ribosome maps from human (EMD-15884), rabbit (EMD-16232), and canine (EMD-3068). D. The hypothesized model of the arrangement of cytoplasmic ribosomes and components associated with the mitochondrial membrane generated by docking atomic models of these components from previous work (Jomaa, Gamerdinger et al. 2022, Gamerdinger, Jia et al. 2023, Ornelas, Bausewein et al. 2023) into our subtomogram average.

**Supplementary Figure 5.**
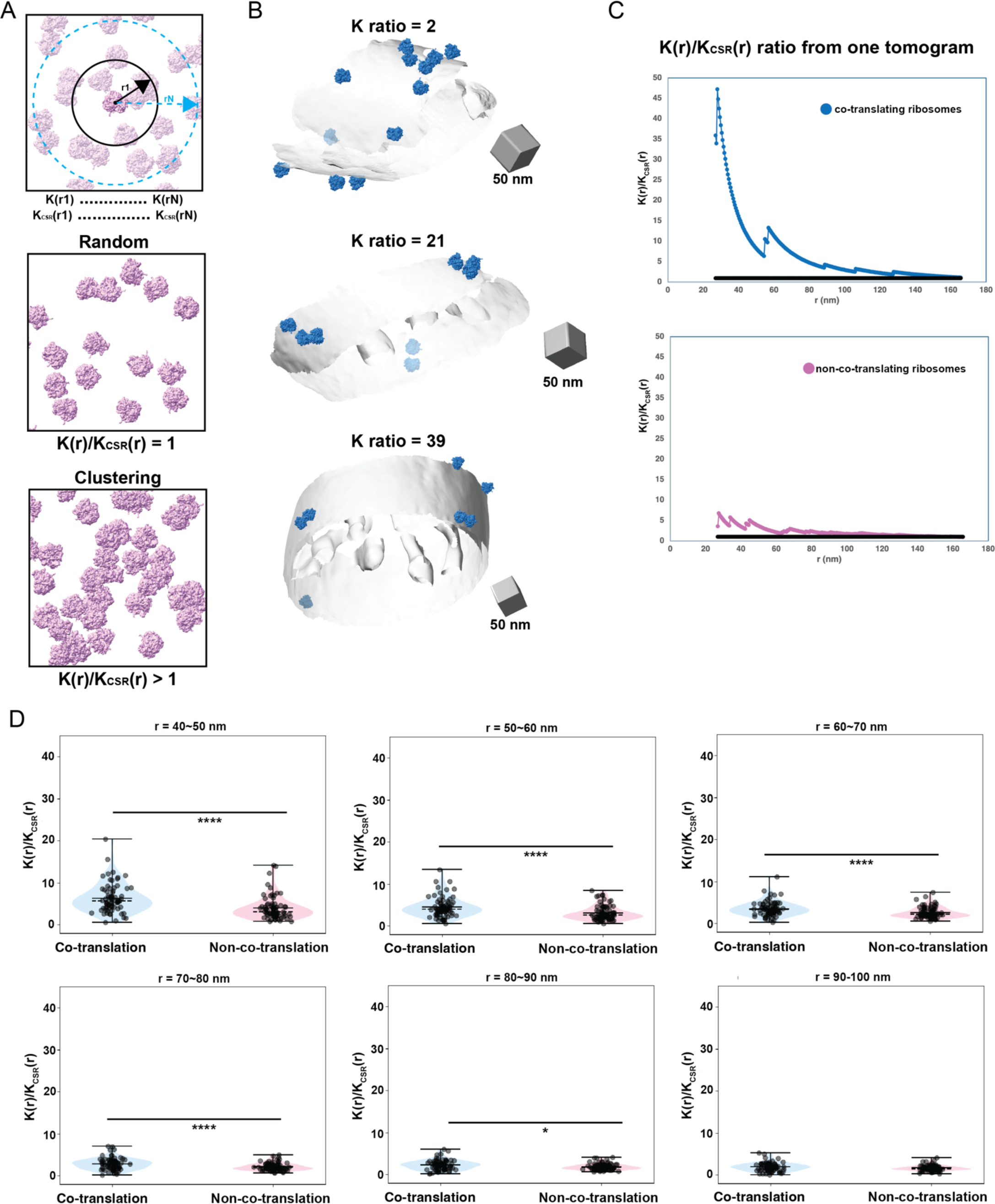
Cytoplasmic ribosomes primed for protein import cluster on the mitochondrial membrane. A. Visual representation of the Ripley’s K function analysis defining the degree of ribosome clustering within a given radius (r) of the indicated ribosome (top panel). The example of spatial organization patterns shows random (middle panel) and clustering (bottom panel) distribution with their corresponding ratio from the analysis. Images are adapted from (Martin-Solana, Diaz-Lopez et al. 2024). B. Representative models of ribosomes oriented for import and membranes from tomograms display different K(r)/K_CSR_(r) ratios. C. Representative plots for the for the K(r)/K_CSR_(r) ratio for mitochondrial-associated cytoplasmic ribosomes oriented for protein import within a range of radius (r) values of 27-166 nm. The black line equals to 1. D. Quantification of the maximum value of K(r)/K_CSR_(r) for each tomogram at the indicated radius intervals for each ribosome class. P values from Mann-Whitney U test are indicated. *P < 0.05; **P < 0.01; ***P < 0.005; ****P < 0.001.

**Supplementary Figure 6.**
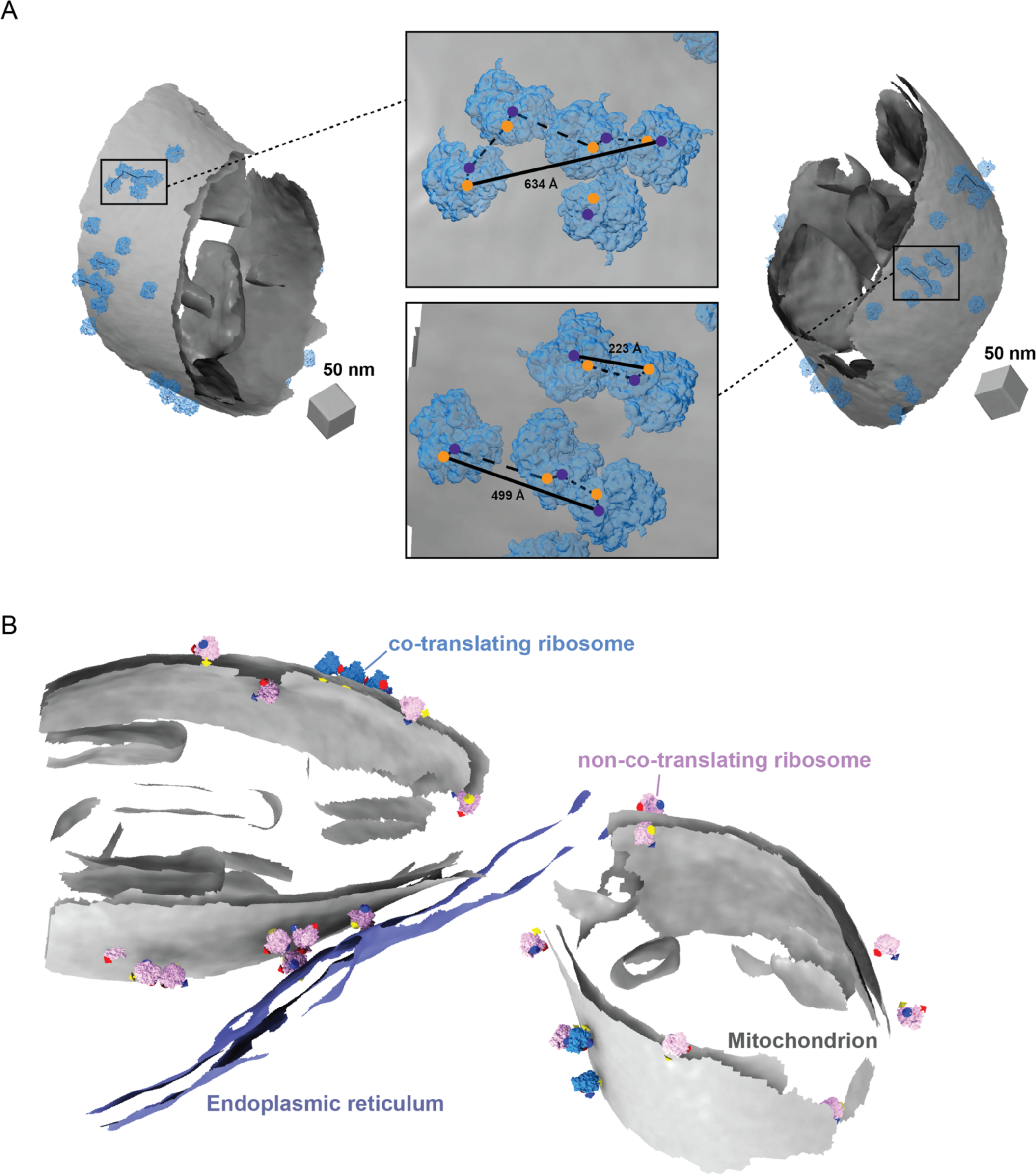
Cytoplasmic ribosomes primed for protein import cluster on the mitochondrial membrane. A. Representative membrane surface reconstructions of mitochondria (gray) with ribosomes oriented for import relative to the OMM (blue). Insets show zoomed-in boxed regions of the ribosome models with circle overlays demarking the location of the 3’ mRNA entry (blue), the 5’ mRNA exit sites (orange), the possible pathways of interconnecting mRNA (dashed black line), and the calculated end-to-end distance from 5’ to 3’ of each interconnected mRNA (solid black line). B. Membrane surface reconstruction of mitochondria (gray) and endoplasmic reticulum (blue) membranes with corresponding models for co-translating (blue) and non-co-translating ribosomes (pink).

**Supplementary Figure 7.**
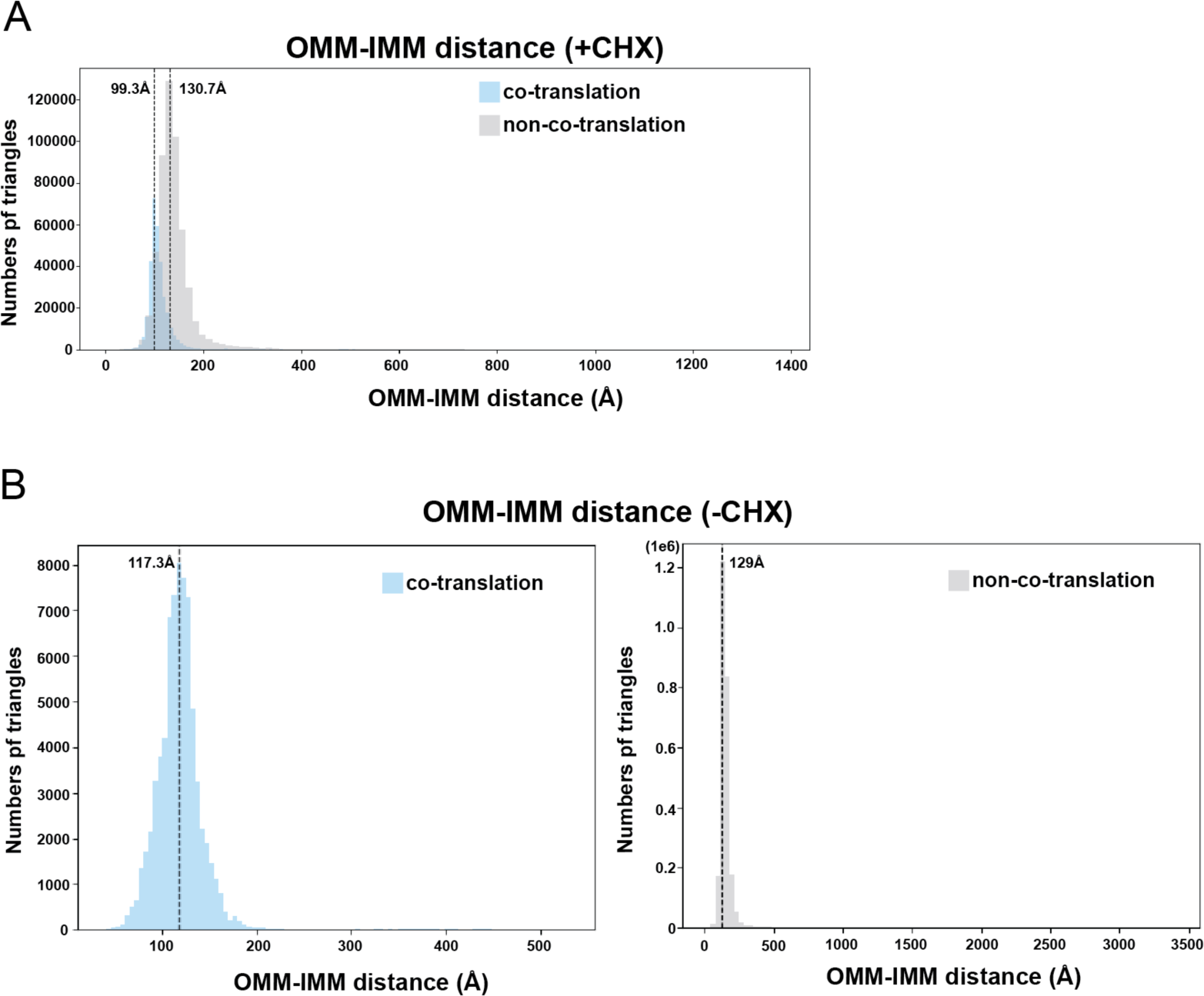
Ribosome-associated protein import alters the local architecture of the outer and inner mitochondrial membranes. A. Combined histogram of IMM-OMM distances of co-translation-associated and non-co-translation-associated patches in *S. cerevisiae* treated with CHX. Dashed vertical lines correspond to peak histogram values of pooled data. B. Histograms of IMM-OMM distances of co-translation-associated and non-co-translation-associated patches in *S. cerevisiae* treated with vehicle (e.g., no CHX). Dashed vertical lines correspond to peak histogram values of pooled data.

**Supplementary Movie 1. Three-dimensional subtomogram average of a cytoplasmic ribosome optimally positioned for protein import on the outer mitochondrial membrane (OMM).**

## REFERENCES

Avendaño-Monsalve, M. C., J. C. Ponce-Rojas and S. Funes (2020). “From cytosol to mitochondria: the beginning of a protein journey.” Biol Chem 401(6-7): 645–661.

Aviram, N. and M. Schuldiner (2017). “Targeting and translocation of proteins to the endoplasmic reticulum at a glance.” J Cell Sci 130(24): 4079–4085.

Barad, B. A., M. Medina, D. Fuentes, R. L. Wiseman and D. A. Grotjahn (2023). “Quantifying organellar ultrastructure in cryo-electron tomography using a surface morphometrics pipeline.” J Cell Biol 222(4).

Becker, T., S. Bhushan, A. Jarasch, J.-P. Armache, S. Funes, F. Jossinet, J. Gumbart, T. Mielke, O. Berninghausen, K. Schulten, E. Westhof, R. Gilmore, E. C. Mandon and R. Beckmann (2009). ”Structure of monomeric yeast and mammalian Sec61 complexes interacting with the translating ribosome.” Science (New York, N.Y.) 326(5958): 1369–1373.

Brandt, F., L. A. Carlson, F. U. Hartl, W. Baumeister and K. Grünewald (2010). “The three-dimensional organization of polyribosomes in intact human cells.” Mol Cell 39(4): 560–569.

Chacinska, A., C. M. Koehler, D. Milenkovic, T. Lithgow and N. Pfanner (2009). “Importing mitochondrial proteins: machineries and mechanisms.” Cell 138(4): 628–644.

Csordás, G., D. Weaver and G. Hajnóczky (2018). “Endoplasmic Reticulum-Mitochondrial Contactology: Structure and Signaling Functions.” Trends Cell Biol 28(7): 523–540.

Deuerling, E., M. Gamerdinger and S. G. Kreft (2019). “Chaperone Interactions at the Ribosome.” Cold Spring Harb Perspect Biol 11(11).

Eisenstein, F., H. Yanagisawa, H. Kashihara, M. Kikkawa, S. Tsukita and R. Danev (2023). “Parallel cryo electron tomography on in situ lamellae.” Nat Methods 20(1): 131–138.

Ermel, U. H., S. M. Arghittu and A. S. Frangakis (2022). “ArtiaX: An electron tomography toolbox for the interactive handling of sub-tomograms in UCSF ChimeraX.” Protein Science 31(12): e4472.

Gamerdinger, M., M. Jia, R. Schloemer, L. Rabl, M. Jaskolowski, K. M. Khakzar, Z. Ulusoy, A. Wallisch, A. Jomaa, G. Hunaeus, A. Scaiola, K. Diederichs, N. Ban and E. Deuerling (2023). “NAC controls cotranslational N-terminal methionine excision in eukaryotes.” Science 380(6651): 1238–1243.

Gemmer, M., M. L. Chaillet, J. van Loenhout, R. Cuevas Arenas, D. Vismpas, M. Gröllers-Mulderij, F. A. Koh, P. Albanese, R. A. Scheltema, S. C. Howes, A. Kotecha, J. Fedry and F. Förster (2023). “Visualization of translation and protein biogenesis at the ER membrane.” Nature 614(7946): 160–167.

George, R., P. Walsh, T. Beddoe and T. Lithgow (2002). “The nascent polypeptide-associated complex (NAC) promotes interaction of ribosomes with the mitochondrial surface in vivo.” FEBS Lett 516(1-3): 213–216.

Gold, V. A., P. Chroscicki, P. Bragoszewski and A. Chacinska (2017). “Visualization of cytosolic ribosomes on the surface of mitochondria by electron cryo-tomography.” EMBO Rep 18(10): 1786–1800.

Gold, V. A., R. Ieva, A. Walter, N. Pfanner, M. van der Laan and W. Kühlbrandt (2014). “Visualizing active membrane protein complexes by electron cryotomography.” Nat Commun 5: 4129.

Guo, T., O. L. Modi, J. Hirano, H. V. Guzman and T. Tsuboi (2022). “Single-chain models illustrate the 3D RNA folding shape during translation.” Biophys Rep (N Y) 2(3): 100065.

Hrabe, T., Y. Chen, S. Pfeffer, L. K. Cuellar, A. V. Mangold and F. Förster (2012). “PyTom: a python-based toolbox for localization of macromolecules in cryo-electron tomograms and subtomogram analysis.” J Struct Biol 178(2): 177–188.

Jaskolowski, M., A. Jomaa, M. Gamerdinger, S. Shrestha, M. Leibundgut, E. Deuerling and N. Ban (2023). “Molecular basis of the TRAP complex function in ER protein biogenesis.” Nat Struct Mol Biol 30(6): 770–777.

Jomaa, A., M. Gamerdinger, H. H. Hsieh, A. Wallisch, V. Chandrasekaran, Z. Ulusoy, A. Scaiola, R. S. Hegde, S. O. Shan, N. Ban and E. Deuerling (2022). “Mechanism of signal sequence handover from NAC to SRP on ribosomes during ER-protein targeting.” Science 375(6583): 839–844.

Kellems, R. E., V. F. Allison and R. A. Butow (1975). “Cytoplasmic type 80S ribosomes associated with yeast mitochondria. IV. Attachment of ribosomes to the outer membrane of isolated mitochondria.” J Cell Biol 65(1): 1–14.

Koch, C., S. Lenhard, M. Räschle, C. Prescianotto-Baschong, A. Spang and J. M. Herrmann (2024). “The ER-SURF pathway uses ER-mitochondria contact sites for protein targeting to mitochondria.” EMBO Rep 25(4): 2071–2096.

Kremer, J. R., D. N. Mastronarde and J. R. McIntosh (1996). “Computer visualization of three-dimensional image data using IMOD.” J Struct Biol 116(1): 71–76.

Lamm, L., S. Zufferey, R. D. Righetto, W. Wietrzynski, K. A. Yamauchi, A. Burt, Y. Liu, H. Zhang, A. Martinez-Sanchez, S. Ziegler, F. Isensee, J. A. Schnabel, B. D. Engel and T. Peng (2024). “MemBrain v2: an end-to-end tool for the analysis of membranes in cryo-electron tomography.” bioRxiv: 2024.2001.2005.574336.

Lesnik, C., Y. Cohen, A. Atir-Lande, M. Schuldiner and Y. Arava (2014). “OM14 is a mitochondrial receptor for cytosolic ribosomes that supports co-translational import into mitochondria.” Nat Commun 5: 5711.

MacKenzie, J. A. and R. M. Payne (2007). “Mitochondrial protein import and human health and disease.” Biochim Biophys Acta 1772(5): 509–523.

Marc, P., A. Margeot, F. Devaux, C. Blugeon, M. Corral-Debrinski and C. Jacq (2002). “Genome-wide analysis of mRNAs targeted to yeast mitochondria.” EMBO Rep 3(2): 159–164.

Martin-Solana, E., I. Diaz-Lopez, Y. Mohamedi, I. Ventoso, J.-J. Fernandez and M. R. Fernandez-Fernandez (2024). “Progressive alterations in polysomal architecture and activation of ribosome stalling relief factors in a mouse model of Huntington’s disease.” Neurobiology of Disease 195: 106488.

Mastronarde, D. N. (1997). “Dual-axis tomography: an approach with alignment methods that preserve resolution.” J Struct Biol 120(3): 343–352.

Maurer, V. J., M. Siggel and J. Kosinski (2024). “PyTME (Python Template Matching Engine): A fast, flexible, and multi-purpose template matching library for cryogenic electron microscopy data.” SoftwareX 25: 101636.

Nyathi, Y. and M. R. Pool (2015). “Analysis of the interplay of protein biogenesis factors at the ribosome exit site reveals new role for NAC.” Journal of Cell Biology 210(2): 287–301.

Ornelas, P., T. Bausewein, J. Martin, N. Morgner, S. Nussberger and W. Kühlbrandt (2023). “Two conformations of the Tom20 preprotein receptor in the TOM holo complex.” Proc Natl Acad Sci U S A 120(34): e2301447120.

Pettersen, E. F., T. D. Goddard, C. C. Huang, E. C. Meng, G. S. Couch, T. I. Croll, J. H. Morris and T. E. Ferrin (2021). “UCSF ChimeraX: Structure visualization for researchers, educators, and developers.” Protein Sci 30(1): 70–82.

Pfanner, N., B. Warscheid and N. Wiedemann (2019). “Mitochondrial proteins: from biogenesis to functional networks.” Nat Rev Mol Cell Biol 20(5): 267–284.

Pfanner, N., N. Wiedemann, C. Meisinger and T. Lithgow (2004). “Assembling the mitochondrial outer membrane.” Nat Struct Mol Biol 11(11): 1044–1048.

Pfeffer, S., F. Brandt, T. Hrabe, S. Lang, M. Eibauer, R. Zimmermann and F. Förster (2012). “Structure and 3D arrangement of endoplasmic reticulum membrane-associated ribosomes.” Structure 20(9): 1508–1518.

Pfeffer, S., L. Burbaum, P. Unverdorben, M. Pech, Y. Chen, R. Zimmermann, R. Beckmann and F. Förster (2015). “Structure of the native Sec61 protein-conducting channel.” Nat Commun 6: 8403.

Saint-Georges, Y., M. Garcia, T. Delaveau, L. Jourdren, S. Le Crom, S. Lemoine, V. Tanty, F. Devaux and C. Jacq (2008). “Yeast mitochondrial biogenesis: a role for the PUF RNA-binding protein Puf3p in mRNA localization.” PLoS One 3(6): e2293.

Schulz, C., A. Schendzielorz and P. Rehling (2015). “Unlocking the presequence import pathway.” Trends Cell Biol 25(5): 265–275.

Suissa, M. and G. Schatz (1982). “Import of proteins into mitochondria. Translatable mRNAs for imported mitochondrial proteins are present in free as well as mitochondria-bound cytoplasmic polysomes.” J Biol Chem 257(21): 13048–13055.

Tegunov, D. and P. Cramer (2019). “Real-time cryo-electron microscopy data preprocessing with Warp.” Nature Methods 16(11): 1146–1152.

Tegunov, D., L. Xue, C. Dienemann, P. Cramer and J. Mahamid (2021). “Multi-particle cryo-EM refinement with M visualizes ribosome-antibiotic complex at 3.5 Å in cells.” Nature Methods 18(2): 186–193.

Tsuboi, T., M. P. Viana, F. Xu, J. Yu, R. Chanchani, X. G. Arceo, E. Tutucci, J. Choi, Y. S. Chen, R. H. Singer, S. M. Rafelski and B. M. Zid (2020). “Mitochondrial volume fraction and translation duration impact mitochondrial mRNA localization and protein synthesis.” Elife 9.

Wang, W., X. Chen, L. Zhang, J. Yi, Q. Ma, J. Yin, W. Zhuo, J. Gu and M. Yang (2020). “Atomic structure of human TOM core complex.” Cell Discovery 6(1): 67.

Wiedemann, N. and N. Pfanner (2017). “Mitochondrial Machineries for Protein Import and Assembly.” Annu Rev Biochem 86: 685–714.

Williams, C. C., C. H. Jan and J. S. Weissman (2014). “Targeting and plasticity of mitochondrial proteins revealed by proximity-specific ribosome profiling.” Science 346(6210): 748–751.

Young, L. N. and E. Villa (2023). “Bringing Structure to Cell Biology with Cryo-Electron Tomography.” Annu Rev Biophys 52: 573–595.

Zivanov, J., J. Otón, Z. Ke, A. von Kügelgen, E. Pyle, K. Qu, D. Morado, D. Castaño-Díez, G. Zanetti, T. A. M. Bharat, J. A. G. Briggs and S. H. W. Scheres (2022). “A Bayesian approach to single-particle electron cryo-tomography in RELION-4.0.” eLife 11: e83724.

